# Parallel Neurodegenerative Phenotypes in Sporadic Parkinson’s Disease Fibroblasts and Midbrain Dopamine Neurons

**DOI:** 10.1101/2023.02.10.527867

**Authors:** MJ Corenblum, A McRobbie-Johnson, E Carruth, K Bernard, M Luo, LJ Mandarino, S Peterson, D Billheimer, T Maley, ED Eggers, L Madhavan

**Affiliations:** Department of Neurology, University of Arizona, Tucson, AZ; Physiological Sciences Graduate Program, University of Arizona, Tucson, AZ; Physiology Undergraduate Program, University of Arizona, Tucson, AZ; Department of Medicine, University of Arizona, Tucson, AZ; Statistical Consulting Lab, University of Arizona, Tucson, AZ; Departments of Physiology and Biomedical Engineering, University of Arizona, Tucson, AZ; Evelyn F McKnight Brain Institute and BIO5 Institute, University of Arizona, Tucson, AZ

## Abstract

Understanding the mechanisms causing Parkinson’s disease (PD) is vital to the development of much needed early diagnostics and therapeutics for this debilitating condition. Here, we report cellular and molecular alterations in skin fibroblasts of late-onset sporadic PD subjects, that were recapitulated in matched induced pluripotent stem cell (iPSC)-derived midbrain dopamine (DA) neurons, reprogrammed from the same fibroblasts. Specific changes in growth, morphology, reactive oxygen species levels, mitochondrial function, and autophagy, were seen in both the PD fibroblasts and DA neurons, as compared to their respective controls. Additionally, significant alterations in alpha synuclein expression and electrical activity were also noted in the PD DA neurons. Interestingly, although the fibroblast and neuronal phenotypes were similar to each other, they also differed in their nature and scale. Furthermore, statistical analysis revealed novel associations between various clinical measures of the PD subjects and the different fibroblast and neuronal data. In essence, these findings encapsulate spontaneous, in-tandem, disease-related phenotypes in both sporadic PD fibroblasts and iPSC-based DA neurons, from the same patient, and generates an innovative model to investigate PD mechanisms with a view towards rational disease stratification and precision treatments.

## INTRODUCTION

Parkinson’s disease (PD) is a relentlessly progressive neurodegenerative disorder characterized by incapacitating motor deficits and neurocognitive decline over the course of aging – but it has no cure (Bloem *et al*., 2021; Dorsey *et al*., 2018; Teves *et al*., 2017). Key to the discovery of effective therapeutics will be an improved understanding of the mechanisms underlying PD etiopathology, and the identification of biomarkers to diagnose the disease at early stages (Poewe *et al*., 2017; Tolosa *et al*., 2021). However, the lack of appropriate model systems that capture the complexity of human PD has hindered progress in this regard.

Although PD research has largely focused on the central nervous system, specifically on the loss of midbrain dopaminergic (DA) neurons and associated motor dysfunction, recently there has been an increasing recognition of the features of PD that are not related to nigrostriatal dopamine deficiency. In fact, PD has emerged as a multisystem disease with numerous clinical manifestations that relate to pathologic changes in widespread regions of both the central and peripheral nervous systems (Lim *et al*., 2009; Poewe, 2010; Shannon, 2007). These clinical features include many non-motor symptoms such as skin sensory and olfactory abnormalities, sleep disturbances, pain, and autonomic dysfunction, in addition to the classic motor symptoms. In terms of its development, PD is understood to originate from complex interactions between several genetic and environmental factors during aging (Poewe *et al*., 2017). Although these gene-environment interactions are not fully understood, studies have demonstrated that converging impairments in mitochondrial function, redox balance, and protein quality control are crucial processes mediating cellular death and dysfunction in PD (Johnson *et al*., 2019; Malkus *et al*., 2009).

In this study, we utilized human skin fibroblasts as well as iPSC-derived dopaminergic neurons (iPSC-DAN), as a two-tiered approach, to examine the cellular and molecular changes defining idiopathic PD. Skin fibroblasts are peripheral cells shown to be valuable in modeling PD as they can express growth, mitochondrial and metabolic alterations relevant to the disease (Carling *et al*., 2020; Galvagnion *et al*., 2022; Mortiboys *et al*., 2008; Teves *et al*., 2017). Importantly, these cells are known to maintain the age and epigenetic signatures of the patient, thus capturing the aging and environmental history, which are the biggest risk factors for PD (Auburger *et al*., 2012; Ivanov *et al*., 2016; Teves *et al*., 2017). In contrast, due to reprogramming, iPSCs are known to revert to a ‘younger state’ mostly devoid of aging markers (Aversano *et al*., 2022; Mertens *et al*., 2018). Thus, iPSCs and their neural progeny, such as DA neurons, may mainly convey mechanisms of early neural vulnerabilities in PD such as the influences of potential genetic variants. Together these two models reflect both early and age-relevant aspects of PD. They also synergistically present peripheral as well as more central (brain related) aspects of the disease. Thus, we propose that human skin fibroblasts and iPSC-derived neural cell types obtained from the same patient may provide a powerful means to investigate PD biology more holistically.

From this perspective, this study assessed a range of processes including growth and morphology, redox status, mitochondrial structure and function, autophagy, alpha synuclein (aSyn) expression and neuronal electrophysiology in the human fibroblasts and iPSC-DAN. Basically, we found distinct changes that differentiated PD fibroblasts, from healthy age- and sex-matched controls. Moreover, these features were recapitulated in midbrain DA neurons, generated from the same fibroblasts, providing ‘neural correlates’ of the fibroblast changes. Interestingly, although several phenotypes in the fibroblasts and DA neurons mirrored each other, they also diverged with respect to their magnitude and precise characteristics. Intriguing correlations between the cellular phenotypes and the patient’s clinical features were also observed. Fundamentally, our study discovers spontaneous disease-related phenotypes expressed in concert in both sporadic PD patient fibroblasts and iPSC-DAN, generating a unique system to deeply study PD development and progression from both peripheral and central mechanistic perspectives.

## METHODS

### Skin Fibroblast and iPSC Lines

Skin fibroblast and iPSC lines of sporadic PD and healthy control subjects were mainly obtained from the Parkinson’s Progression Markers Initiative (PPMI) biorepository at Indiana University (supported by the Michael J. Fox Foundation and it corporate sponsors) with some lines from the Madhavan lab cell bank. All the chosen fibroblast lines were originally generated from upper arm skin biopsies collected under relevant Institutional Review Board approvals. 9 control (C1-C9) and 8 PD (PD1-PD8) lines, which were age-matched, and contained 4-5 male and 4 female samples in each group from passages 4-5 were utilized for the study. The detailed clinical and demographic information related to these cell lines are described in **Table 1**. Induced pluripotent stem cells (iPSCs) derived from a subset of the fibroblast lines were also studied. Specifically, already generated and quality control checked iPSC lines from C1, C6, C8, C9, PD1, PD6, PD4 and PD8 fibroblasts, from the same sources mentioned above, were used. The iPSC lines were age-matched and contained 2 male and 2 female lines in each group.

**Table 1.** Demographics & Clinical Data.

| Line ID | Group | Sex | Age at diagnosis (years) | Age at biopsy (years) | Disease Duration (years) | UPDRS3 (ON) | UPDRS TOT (ON) | UPDRS3 | UPDRS TOT | Cog State | NPI-Cog | REM-Cat | A-Syn | PD Med Use |
| --- | --- | --- | --- | --- | --- | --- | --- | --- | --- | --- | --- | --- | --- | --- |
| CON1 | Control | M |  | 75 |  |  |  |  |  |  |  |  |  |  |
| CON2 | Control | M |  | 80 |  |  |  |  |  |  |  |  |  |  |
| CON3 | Control | M |  | 74 |  |  |  |  |  |  |  |  |  |  |
| CON4 | Control | M |  | 59 |  |  |  |  |  |  |  |  |  |  |
| CON5 | Control | F |  | 66 |  |  |  |  |  |  |  |  |  |  |
| CON6 | Control | F |  | 68 |  |  |  |  |  |  |  |  |  |  |
| CON7 | Control | F |  | 59 |  |  |  |  |  |  |  |  |  |  |
| CON8 | Control | F |  | 61.5 |  |  |  |  |  |  |  |  |  |  |
| CON9 | Control | M |  | 44.5 |  |  |  |  |  |  |  |  |  |  |
|  | Ratio F:M/ Mean | 4:5 |  | 65.2 |  |  |  |  |  |  |  |  |  |  |
| PD1 | PD | M | 65.2 | 69.2 | 4 | 43 | 51 | 25 | 33 | Normal | 0-Normal | 1-No RBD | 471.5 | 2-DA Agonist |
| PD2 | PD | M | 68.8 | 71.3 | 2.5 | 26 | 55 | 35 | 64 | Normal | 1-Slight | 1-RBD | 2361.8 | 4-Ldopa + Other |
| PD3 | PD | M | 48.1 | 50.7 | 2.6 | 28 | 35 | 28 | 35 | Normal | 0-Normal | 0-No RBD | 998.1 | 3-Other |
| PD4 | PD | M | 75.5 | 77.8 | 2.3 | 21 | 46 | 27 | 52 | Normal | 1-Slight | 0-No RBD | 1313.2 | 1-Ldopa |
| PD5 | PD | F | 67.4 | 70.7 | 2.3 | 10 | 13 | 10 | 13 | Normal | 0-Normal | 0-No RBD | 972.1 | 0-Unmedicated |
| PD6 | PD | F | 61.6 | 64.5 | 2.9 | 12 | 30 | 12 | 30 | Normal | 0-Normal | 0-No RBD | 1313.6 | 2-DA Agonist |
| PD7 | PD | F | 81.8 | 84.8 | 3 | 15 | 27 | 15 | 27 | Normal | 0-Normal | 1-RBD | 1933.9 | 1-LDopa |
| PD8 | PD | F | 53.1 | 55.3 | 2.2 | 26 | 35 | 26 | 35 | Normal | 0-Normal | 0-No RBD | 804.5 | 3-Other |
|  | Ratio F:M/ Mean | 4:4 | 65.2 | 68 | 2.725 | 24 | 37 | 22 | 35.5 |  |  |  |  |  |

NA, not available; UPDRS, Unified Parkinson's Disease Rating Scale Part 3 (on & off medications); UPDRS TOT, Unified Parkinson's Disease Rating Scale Total (on & off medications); Cog state; Diagnosis of cognitive state (1-normal, 2-MCI, 3-Dementia); NPI-Cog, Neuropsychiatric Inventory Score of Cognitive Impairment (0-Normal, 1-slight, 2-mild, 3-moderate, 4-severe); REM-Cat, REM Sleep Behavior Disorder Categorization (0-No RBD, 1-RBD); A-Syn, alpha-synuclein levels; PD Medication Use, Medications taken by the patient for PD symptoms (0-unmedicated, 1-Ldopa, 2-DA agonist, 3-Other, 4-Ldopa + other, 5- DA agonist + other).

### Cell Culture

#### Fibroblasts

Fibroblasts were grown in Dulbecco’s Modified Eagle’s Medium (DMEM) (Thermofisher Scientific, Cambridge, MA), supplemented with 1X Non-Essential Amino Acid (NEAA) (Thermofisher Scientific, Cambridge, MA), 10% Fetal Bovine Serum (FBS) (Atlanta Biologicals, Flowery Branch, GA), and 1% Anti-mycotic Anti-biotic (Anti-Anti) (Gibco, Waltham, MA) at 5% CO_2_ and 37°C. Fibroblasts from passage 7 to10 were used, and for all experiments, the passage numbers were kept consistent within groups to avoid cell replication related biases. Cells were plated and used for the different assays when they reached ~75% confluence in culture. All fibroblast lines were grown in parallel and assessed in at least triplicate for the experiments.

#### Induced pluripotent stem cells (iPSCs)

iPSCs were maintained on Matrigel hESC-Qualified Matrix (Corning, Corning NY) in mTeSR Plus (Stemcell Technologies, Vancouver BC). Colonies were routinely clump passaged using enzyme-free Gentle Cell Dissociation Reagent (Stemcell Technologies).

**Figure 1:**
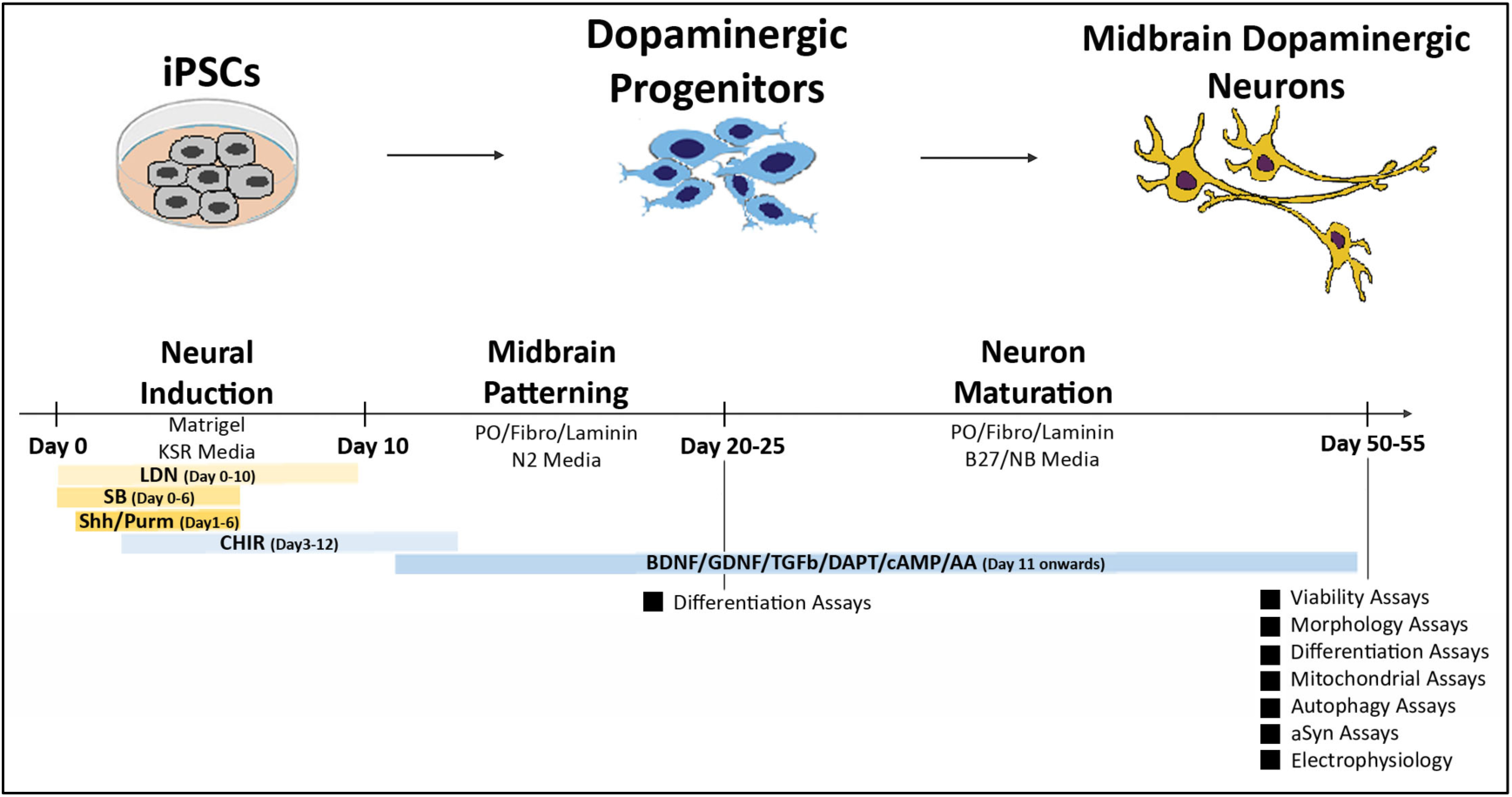

For deriving midbrain DA neurons, a modified floor plate-based differentiation method involving dual SMAD inhibition was used (Chambers *et al*., 2009; Kriks *et al*., 2011). This method involves initiating the neural patterning of the iPSCs via inhibitors of the SMAD signaling pathway [LDN 193189 (100nM, Tocris, Minneapolis MN), SB 431542 (10μM, Tocris), and specification towards a ventral midbrain DA neuron fate using the morphogens sonic hedgehog (SHH C25II N-Terminus 100ng/mL, R&D Systems, Minneapolis MN), Purmorphamine (2μM, Tocris), and CHIR 99021 (3μM, Tocris). This is followed by the stimulation of neuronal maturation through the application of ascorbic acid, brain derived neurotrophic factor (BDNF), glial cell-line derived neurotrophic factor (GDNF), Transforming growth factor type β3 (TGF-b3), DAPT, L-Ascorbic acid (AA), and cyclic AMP (cAMP) on Poly-L-Ornithine/Fibronectin/Laminin coated dishes [BDNF: 20ng/mL, Peprotech, Cranberry NJ, GDNF: 20ng/mL, Peprotech, TGF-b3: 1ng/mL, Peprotech, DAPT: 10uM, Biogems, Westlake Village CA, AA: 200uM, Sigma-Aldrich, St. Louis MO, and cAMP: 0.5mM, Sigma-Aldrich, Poly-L-Ornithine: mw30,000-70,000; 15μg/ml, Sigma-Aldrich, Fibronectin: 4μg/ml, Corning, Laminin: 6ug/mL, Sigma]. Cells were assessed after 25 (dopaminergic progenitor stage) and 50-55 (more mature DA neurons) days of differentiation, when they were subjected to a variety of assays. All iPSC lines were differentiated in parallel and assessed in at least triplicate for all experiments. The schematic above gives a broad overview of the iPSC differentiation and characterization process. A more detailed methodology has been provided in the **Supplementary Methods** section.

### Cell Viability

Cellular viability was determined utilizing a MTT [3-(4,5-dimethylthiazol-2-yl)-2,5-diphenyltetrazolium bromide] assay (Invitrogen, Waltham MA), as described previously (Kumar *et al*., 2018; Teves *et al*., 2017). Fibroblasts and iPSC-DAN were detached and processed into single cell suspensions and plated onto 96-well clear flat-bottom plates at 10,000 cells/well and 80,000 cells/well respectively (at least 4 replicates per experimental run). Cell negative and assay negative controls were also used. Briefly, cells were washed once with PBS and placed in phenol-free culture medium. Experimental wells and cell negative wells were then treated with 12mM MTT (prepared as described in assay manual) at 37°C for 2hrs. Assay negative wells received PBS only and were incubated in the same manner. A portion of the MTT solution was removed from each well and Dimethyl sulfoxide (DMSO; Sigma-Aldrich) added to solubilize the generated crystals. Absorbance was read on a standard plate reader (Tecan Infinite 200 Pro Plate Reader, Tecan, Zurich, Switzerland) at 540nm.

### Fibroblast Growth and Morphology

The growth and morphology of the fibroblasts were assessed when they reached 75% confluence (right before they are ready to be passaged) as described previously (Teves *et al*., 2017). Three variables pertaining to cellular growth were measured across 4 passages: (1) duration (in days) it took for the cells to reach 75% confluency; (2) total cell count at 75% confluency, and (3) population doubling time [PDT = 3.32 × [Log (number of cells at 75% confluence) − Log (number of cells plated) + number of cells/ cm^2^].

For morphological analysis, the fibroblasts were plated at 40,000 cells/well on poly-D-lysine (0.1mg/mL) coated glass coverslips in 24-well plates. The cells were fixed using 4% paraformaldehyde (PFA, Electron Microscopy Sciences, Hatfield, PA) for 20min at room temperature. After washing with 1X Phosphate-buffered saline (PBS: Thermofisher Scientific, Cambridge, MA), the fibroblasts were treated with 0.1% Triton-X-100 (Sigma-Aldrich, Darmstadt, Germany) in PBS for 5min. Cells were then stained with 1.65uM Alexa Fluor 488 Phalloidin (Invitrogen, Waltham, MA) for 20min at room temperature. Images were subsequently obtained from 6 random fields per coverslip using a Zeiss Axio Imager M2 Upright Fluorescent Microscope (Zeiss, Jenna, Germany). Images were taken at 40X with exposure conditions standardized across all cell lines. Afterwards, the single-channel images were processed in CellProfiler open-source software (Broad Institute at MIT, Cambridge, MA). An analysis pipeline was created to correctly identify individual nuclei and cell perimeter/outlines. Using this pipeline, numerous parameters of interest were analyzed including the density (neighbors), size (area and perimeter) and shape (form factor and eccentricity) of cells for all lines.

### Immunocytochemistry

Fibroblasts were enzymatically dissociated at ~75% confluence and 20,000 cells plated on glass coverslips coated with 0.1mg/mL Poly-D-Lysine (Sigma-Aldrich, Darmstadt, Germany) and placed in 24-well plates. iPSC-DAN were dissociated and replated on glass coverslips at 150,000 cells per coverslip or 100,000 cells per well of an 8-well chamber slide, both coated with Poly-L-Ornithine (15ug/mL Sigma), Fibronectin (4ug/mL Corning), Laminin (6ug/mL Sigma). All cells were fixed for 20min at room temperature (RT) with 4% PFA. For immunocytochemistry, cells were blocked for 2hrs at room temperature [in 2% normal goat serum, 1% Bovine Serum Albumin, 0.4% Triton-X-100 in PBS] and incubated in primary antibody overnight at 4°C in a humidity chamber. Primary antibodies were detected in a 2-hour incubation at RT with secondary antibodies coupled to fluorochromes Alexa Fluor 488, 594, 647 (Life Technologies-Molecular Probes, Grand Island, NY) or a biotinylated secondary (Vector Laboratories, Newark CA) followed by a Streptavidin-tagged Alexa Fluor, and counterstained with 4’,6’-diamidino-2-phenylindole, dihydrochloride (DAPI, Life Technologies). Control conditions constituted the deletion of the primary antibody or secondary antibody and the inclusion of relevant isotype specific antibodies and sera instead of the omitted antibodies. Primary antibodies used were as follows: ATPIF1 (1:350); Tuj1 (1:200); TH (1:2000); LMX1A (1:250); FoxA2 (1:200); Girk2 (1:1000); Map2 (1:8000); aSyn (1:400). More details can be found in the table in the **Supplementary Methods** section.

### Microscopy and Cell counts

Fluorescence analysis was performed using a Zeiss LSM880 Inverted confocal microscope (Zeiss, Jena, Germany). Z sectioning was performed at 1–2μm intervals in order to verify the co-localization of markers. Image extraction and analysis was conducted via the Zen Blue software. A Zeiss Axio-Imager M2 microscope connected to an Axiocam MRm digital camera and Microlucida software (v2019.1.3, MBF Bioscience) was used for epifluorescence microscopy. A Zeiss Axio Observer A1 inverted microscope with Axiocam MRc camera and AxioVision software was used for phase contrast microscopy. Quantitative analyses were conducted by a person blinded to the groups and lines.

#### Cell counts

For the TH and TH/Girk2 cell counts 5 random fluorescence images were taken at 63X magnification of cells in 8-well chamber slides (duplicate wells/line), and the number of positively stained cells was elucidated and expressed as percent of Dapi. For quantification of the aSyn mean intensity signal, 5 random fluorescence images were taken at 63X magnification of cells in 8-well chamber slides (duplicate wells/line) keeping the exposure and gain settings consistent among all lines and images. Individual images were loaded into Cell Profiler (Broad Institute) and a set pipeline was run which identified each cell’s nucleus and quantified the intensity of its surrounding aSyn staining on a pixel-by-pixel basis. Quantification of neuronal soma size and number of neurites was conducted using NIH Image J software. 5 random epifluorescence images were taken at 63X magnification of Map2 staining of cells in 8-well chamber slides (duplicate wells/line). Images were loaded into Image J and a freehand selection tool was utilized to precisely trace each cell’s soma and the area was measured in square pixels and number of primary neurites was enumerated.

#### Mitochondrial network analysis

Confocal images were acquired of the ATPIF1 immunostaining using a 63X objective (3 coverslips and 15 images/line). Z-stacks were acquired at 0.38uM intervals and compressed to a 2D maximum intensity projection image using the ImageJ software (NIH, v1 53c).

*Skeleton Analysis:* From single channel images, a single cell was outlined, and the outside cleared such that only one cell remained per frame. Images were converted to binary, and total area was calculated using the measure feature. Morphological features were quantified using an objective and computer-aided skeleton analysis method previously described in detail (Young and Morrison, 2018). After thresholding, a series of ImageJ plugins were consistently applied across all images to ensure adequate visualization of cell process before the conversion to binary and skeletonized images. The skeletonized representations of original photomicrographs were used for data collection of morphology parameters using the AnalyzeSkeleton (2D/3D) plugin (Arganda-Carreras *et al*., 2010). These features were then divided by the total area of each cell, in order to normalize to the total size of each cell.

*Fractal and Lacunarity Analysis:* Single channel images were converted to binary, and then run through the ImageJ plugin, FracLac. For fractal analysis the program was set up so that NumG = 5. All binary images were run through the program using the batch feature. D_B_, Lacunarity, and Density of Foreground Pixels were selected from the resulting data table.

### Autophagy flux assays

To measure autophagic flux, confluent wells in a 6-well plate were incubated with 20mM NH_4_Cl (Sigma-Aldrich, St Louis MO) and 300μM leupeptin (Sigma-Aldrich) for 4hrs in standard culture medium with serum or serum-free conditions before protein samples were collected as below and western blotting was performed (Teves *et al*., 2017).

### Western Blotting

To isolate protein, fibroblasts and DA neurons were trypsinized, washed in PBS, and resuspended in RIPA buffer (Sigma-Aldrich, St. Louis MO) containing Protease Inhibitor Cocktail (Sigma-Aldrich). After 1hr incubation in RIPA on ice, cells were sonicated and centrifuged at 4°C for 30min at 15,000 × *g*. The supernatant containing the soluble protein was removed, quantified by the Lowry method, and stored at −20°C. To detect LC3 and p62, samples from Control and PD fibroblasts and DA neurons were run on a 12% acrylamide gel and transferred to a PVDF membrane using a wet transfer system. For aSyn, protein samples were run on a 12.5% acrylamide gel and subsequently transferred to a PVDF membrane. After 1hr of incubation with blocking solution [0.1M tris buffered saline (TBS) with 1% bovine serum albumin (BSA, ThermoFisher Scientific, Waltham MA) and 5% dry milk], primary and secondary antibodies were applied. Specifically, membranes were incubated overnight in primary antibodies targeting LC3 (1:2000) and p62 (1:2000) and aSyn (1:500) diluted in blocking solution with 0.1% Tween-20 (Sigma-Aldrich). The next day, after washing in 0.1M TBS with 0.1% Tween-20, membranes were incubated in appropriate horseradish peroxidase secondary antibodies for 1hr and developed using SuperSignal west Femto ECL substrate (ThermoFisher) and scanned on an Azure 500 (Azure Biosystems, Dublin CA).

### Transmission Electron Microscopy

Fibroblasts were plated at 1×10^6^ cells per 10mm petri dish and allowed to grow to confluence. iPSC-DAN were plated in 6-well plates and allowed to differentiate and mature to Day 50. The cells were fixed in-situ with 2.5% glutaraldehyde in 0.1M piperazine-N,N’-bis (1-ethanesulfonic acid) (Glut-PIPES buffer) for 1hr at room temperature, washed 3 times with 0.1M PBS pH 7.4, and scraped off well bottoms. Then the cells were transferred to a 1.5mL microcentrifuge tube and centrifuged at 1000 × g. Afterwards, the cell pellets were post-fixed with 1% osmium tetroxide in 0.1M PBS for 30min, resuspended and washed in the same buffer and in distilled/deionized water. Subsequently, the cell pellets were dehydrated through a series of ethanol and acetonitrile and infiltrated with 1:1 acetonitrile/ Spurr resin overnight. The next day pellets were embedded in 3 changes of Spurr resin 60min each and allowed to polymerize in Beem embedding capsules for 36hrs at 60°C. Thin sections (90nm) were obtained on a RMC PowerTome XL 9 ultramicrotome onto uncoated 150 mesh copper grids and stained with 2% uranyl acetate and lead citrate. Images were acquired using a Tecnai G2 Spirit BioTwin transmission electron microscope (FEI, Hillsboro, OR USA) with a side mounted XR41 AMT 4 Mpix digital camera, operated at 100kV. Morphometric measurements were conducted in digital images and the number of autophagic vacuoles or distorted mitochondria per cell profile (10-15 cell profiles) across a minimum of 10 micrographs per line were counted using standard criteria as described before (Anandhan *et al*., 2021; Teves *et al*., 2017).

### Reactive Oxygen Species (ROS) measurements

ROS levels were assessed using the DCFDA/H2DCFDA Cellular ROS Assay Kit (Abcam, Waltham MA), as describe before (Oyama *et al*., 1994; Teves *et al*., 2017). Fibroblasts and iPSC-DAN neurons were detached and processed into single cell suspensions and plated onto 96-well black-walled flat-bottom plates at 10,000 cells/well and 80,000 cells/well, respectively, to include at least 4 replicates per experimental run. During the assay, following manufacturer’s instructions, culture media was removed and 25uM DCFDA solution was added. Cell negative and assay negative controls were also used. Cells were incubated for 45min at 37°C in DCFDA solution, after which they were washed in PBS, and fluorescence measured at 485Ex/535Em on a standard plate reader (Tecan Infinite 200 Pro Plate Reader, Tecan, Zurich, Switzerland). Values were normalized to protein expression using a standard Bradford assay.

### Seahorse Mito Stress Test

Mitochondrial function was probed utilizing a Seahorse XF Cell Mito Stress Test kit (Agilent Technologies, Santa Clara CA) (Teves *et al*., 2017). Fibroblasts were plated in a Seahorse XFe96 V3 PS plate at an optimized seeding density of 20,000 cells per well, and each consisted of at least 8 replicates per experimental run. iPSC-DAN were plated in a Seahorse XF24 V7 PS plate at an optimized seeding density of 100,000 cells per well and consisted of at least 5 replicates per line. Cells were cultured in 5% CO_2_ and 37°C to allow them to adhere and grow in the plate. Seahorse XF base medium enriched with 10mM glucose, 2mM L-glutamine and 1mM sodium pyruvate was equilibrated to 37°C with an adjusted pH of 7.35±0.05. All wells were carefully washed 3 times with the Seahorse XF base medium and then incubated for 1hr in assay media in a 37°C CO_2_-free incubator before the Mito Stress test was conducted. Successive administration of Oligomycin (Oligo), Carbonyl cyanide p-trifluoro-methoxyphenyl hydrazone (FCCP), and Rotenone/ Antimycin (R/A), all modulators of respiration, were performed in the Seahorse XF Flux Analyzer (Fibroblasts: 1µM Oligo, 2µM FCCP, 1µM/1uM R/A. DA neurons: 1µM Oligo, 1µM FCCP, 2µM/2uM R/A. Values were normalized to protein concentration via a standard Bradford assay.

### Electrophysiology assays

iPSC-DAN were plated on individual glass cover slips and kept in culture media at 37°C until the time of experiments. Cells were removed from culture media and immediately used for recording. Extracellular solution used for whole cell recordings was kept at room temperature and contained the following (in mM): 125 NaCl, 2.5 KCl, 1.00 MgCl_2_, 1.25 NaH_2_PO_4_, 20 glucose, 26 NaHCO_3_, and 2 CaCl_2_. The intracellular solution in the recording pipette contained the following (in mM): 120 K D-Gluconate, 25 KCl, 10 HEPES, 10 EGTA, 4 phosphocreatine-Na2, 4Mg-ATP, 2 Na-GTP, 1 CaCl_2,_ and 0.1% sulforhodamine-B dissolved in water and was adjusted to pH 7.2 with CsOH.

Electrodes were pulled from borosilicate glass (World Precision Instruments, Sarasota, FL) using a P97 Flaming/Brown puller (Sutter Instruments, Novato, CA) and had resistances of 5–8 MΩ. Liquid junction potentials of 20mV, calculated with Clampex software (Molecular Devices, Sunnyvale, CA), were corrected before recording. Spontaneous events were sampled at 10kHz and filtered at 6kHz with the four-pole Bessel filter on a MultiClamp 700B patch-clamp amplifier (Molecular Devices) before being digitized with a Digidata 1140 data acquisition system (Molecular Devices) and Clampex software.

iPSC-DAN were considered spontaneously active if they spiked at least once without current steps. Cells were considered inducibly active if they elicited spikes after or during a −30pA or +30pA current step. Non-active cells never exhibited meaningful changes in membrane voltage. Spontaneous cell frequency, peak amplitude, decay tau, inter-event interval, and time to peak were analyzed using Molecular Devices Clampfit threshold search event detector software. Means were analyzed using Sigmaplot software to compute ANOVA SNK post-hoc test. Differences were considered significant when p ≤ 0.05. Membrane capacitance was measured at the time of recording.

### Statistical Analysis

GraphPad Prism 9 software (San Diego, CA) was used for statistical analyses. For comparing two groups, unpaired *t* tests were used. For comparisons between three or more groups, one-way analysis of variance (ANOVA) followed by Tukey’s or Bonferroni’s post-hoc test for multiple comparisons between treatment groups was conducted. When there were two independent variables involved, a two-way ANOVA test with Tukey’s multiple comparisons was used. For the ATPIF1 image analysis, in order to assess the morphological differences between groups while also capturing the variability between cells within the same line, each cell was representative of itself, and images were not averaged across lines. In all cases, differences were accepted as significant at *p* < 0.05. For the statistical correlation data, R version 4.2.1 was used for performing the analysis and graphs. The plots were all created using the R package ggplot2 (https://ggplot2.tidyverse.org). To produce the correlation matrices, Corrplot (https://github.com/taiyun/corrplot) package in R was used. This package calculates coefficients based on the Pearson’s correlation. The exact sample size and specific statistical details of each experiment are provided within the relevant result and legend sections.

## RESULTS

### PD and Control fibroblasts show different growth patterns and morphology

Several fibroblast lines from sporadic PD subjects and age- and sex-matched healthy control individuals were studied. The details of the cell lines and the associated clinical and demographic information is displayed in **Table 1**. First, the growth and morphological characteristics of fibroblasts were examined. All assessments were conducted right before passage when the cells had reached ~75% confluence in culture. At this stage, it was observed that cells within control cultures were large, well ramified, and evenly distributed over the culture surface, as typical of mature fibroblasts. PD fibroblasts on other hand were smaller and more spindle shaped. Additionally, the PD cultures were generally denser and contained closely packed cells aligned along their longitudinal axis (**Fig. 1A, B** - phase contrast images; **Fig. 1C, D** - high magnification fluorescent images of Phalloidin/DAPI stained cells, Phalloidin binds to F-Actin).

**Figure 1:**
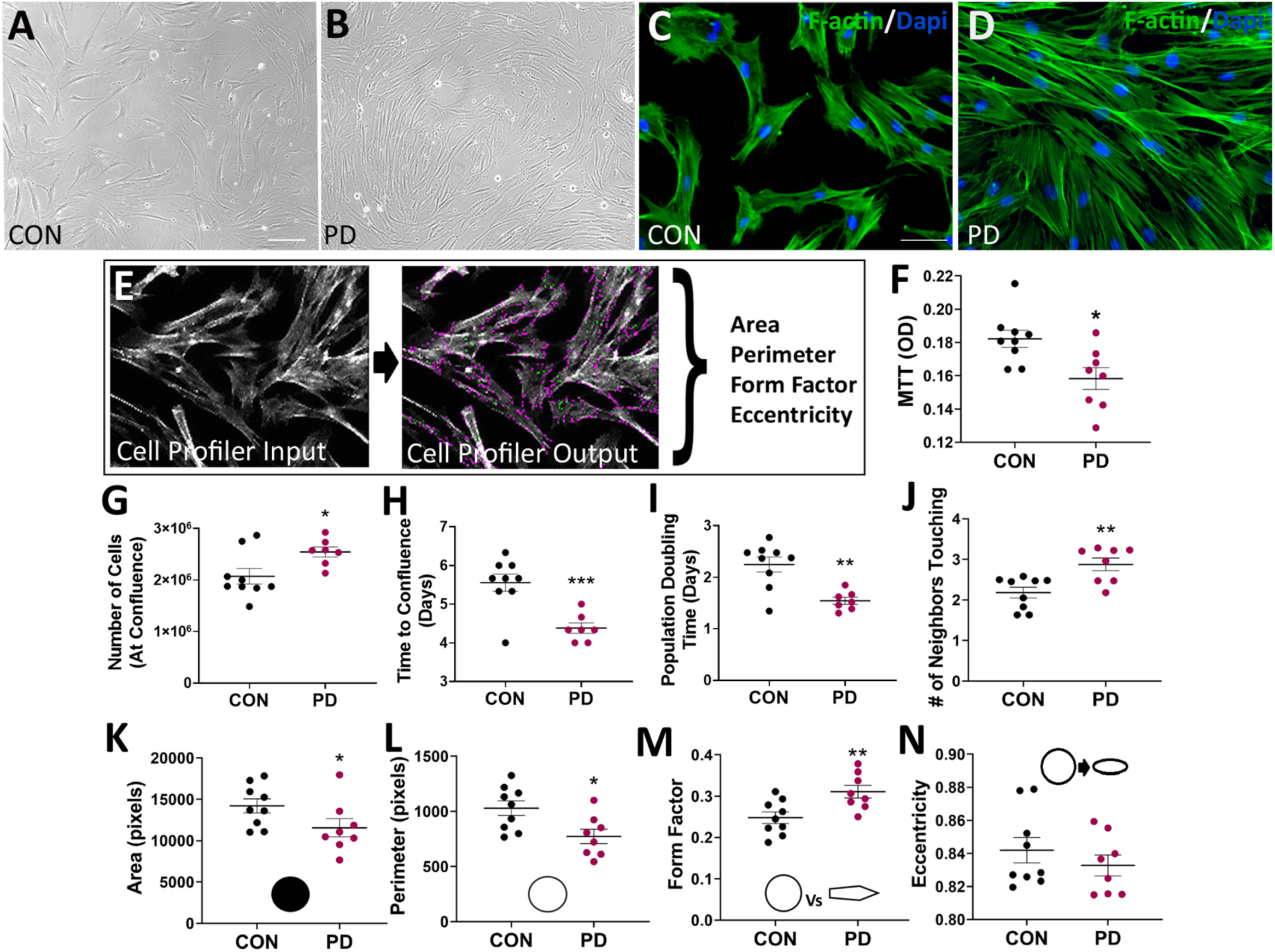
Growth dynamics and morphological properties of the patient-derived fibroblasts. CON and PD fibroblasts exhibited perceptible differences in their spatial growth patterns as depicted in the phase images in (A-B). Staining the cells with Phalloidin (fluorescent tag that binds to F-Actin) allowed to clearly visualize these growth differences and enabled a careful assessment of their morphological properties via NIH Cell Profiler (C, D, I). (H) shows data comparing the cellular viability of PD vs CON fibroblasts using an MTT assay. Comparisons between the number of live cells at 75% confluence, the time taken to reach 75% confluence and the population doubling time are in (E-G). The number of cellular neighbors is shown in (J). Results from shape and size assessments, specifically area (K), perimeter (L), form factor (M), and eccentricity (N), between CON and PD fibroblasts are in (K-N). Scale Bars: (A-B) = 200μm, (C-D) = 100μM. *p<0.05, **p<0.01; Mean ± SEM, Unpaired t-tests, n = 8-9 independent lines/group.

To further investigate these observations, we quantified the total number of cells in the culture flasks at 75% confluence and assessed their growth rate. It was found that there were significantly higher numbers of cells in the PD flasks (**Fig. 1E**; p < 0.05, Unpaired t-test), and the time taken to reach 75% confluence was shorter with the PD cultures (~4.5 days) compared to controls (~5.5 days) (**Fig. 1F**; p < 0.001, Unpaired t-test). Additional analysis indicated that the population doubling time (PDT) of control fibroblasts was significantly faster than that of PD cells (**Fig. 1G**; p < 0.01, Unpaired t-test). Furthermore, PD cultures were found to be denser, with each cell surrounded by more neighboring cells, than controls (**Fig. 1J**; p < 0.01, Unpaired t-test). Interestingly, although more cells were detected in the PD cultures, it was also determined that that PD cells had lower viability rates than control cells (**Fig. 1H**; p < 0.05, Unpaired t-test).

The cellular morphology of the fibroblasts was also assessed. Specifically, the fibroblasts were stained with Phalloidin and DAPI to allow clear demarcations of cell size and shape, and subsequently analyzed via the CellProfiler software (**Fig. 1I**). Cell size (area and perimeter), form factor, and eccentricity were calculated. On average, PD cells had significantly lower area (**Fig. 1K**; p < 0.05, Unpaired t-test) and perimeter (**Fig. 1L**; p < 0.05, Unpaired t-test), indicating that they were smaller compared to control cells. With regards to shape, the eccentricity (a measure of cell elongation) and form factor (a measure of roundness) of the fibroblasts were measured. It was seen that although PD and control cells were both elongated (**Fig. 1N**), PD fibroblasts were significantly more defined (less ramified and rounder) than control fibroblasts (**Fig. 1M**; p < 0.01, Unpaired t-test). These data demonstrate that PD fibroblasts distinctly differed in terms of both their growth and morphological characteristics from controls.

### PD Fibroblasts exhibit impaired mitochondrial structure and function

We assessed mitochondrial function in the fibroblasts using the Seahorse Mito Stress test (**Fig. 2**). The primary function of mitochondria is oxidative phosphorylation (OXPHOS) to produce the majority of ATP. Apart from their metabolic function, the mitochondria also play a crucial role in reactive oxygen species (ROS) generation (Murphy, 2009; Venditti *et al*., 2013). The Seahorse Mito Stress Test estimates key parameters of mitochondrial function by directly measuring the oxygen consumption rate (OCR) of cells after the addition of specific modulators of respiration (Gu *et al*., 2021; Teves *et al*., 2017). **Figure 2A** illustrates the complexes of the Electron Transport Chain (ETC) and indicates the target of action of the modulators used in the Seahorse Mito Stress Test which then reveals the key parameters of mitochondrial function, some of which are depicted in **Figure 2C**.

**Figure 2:**
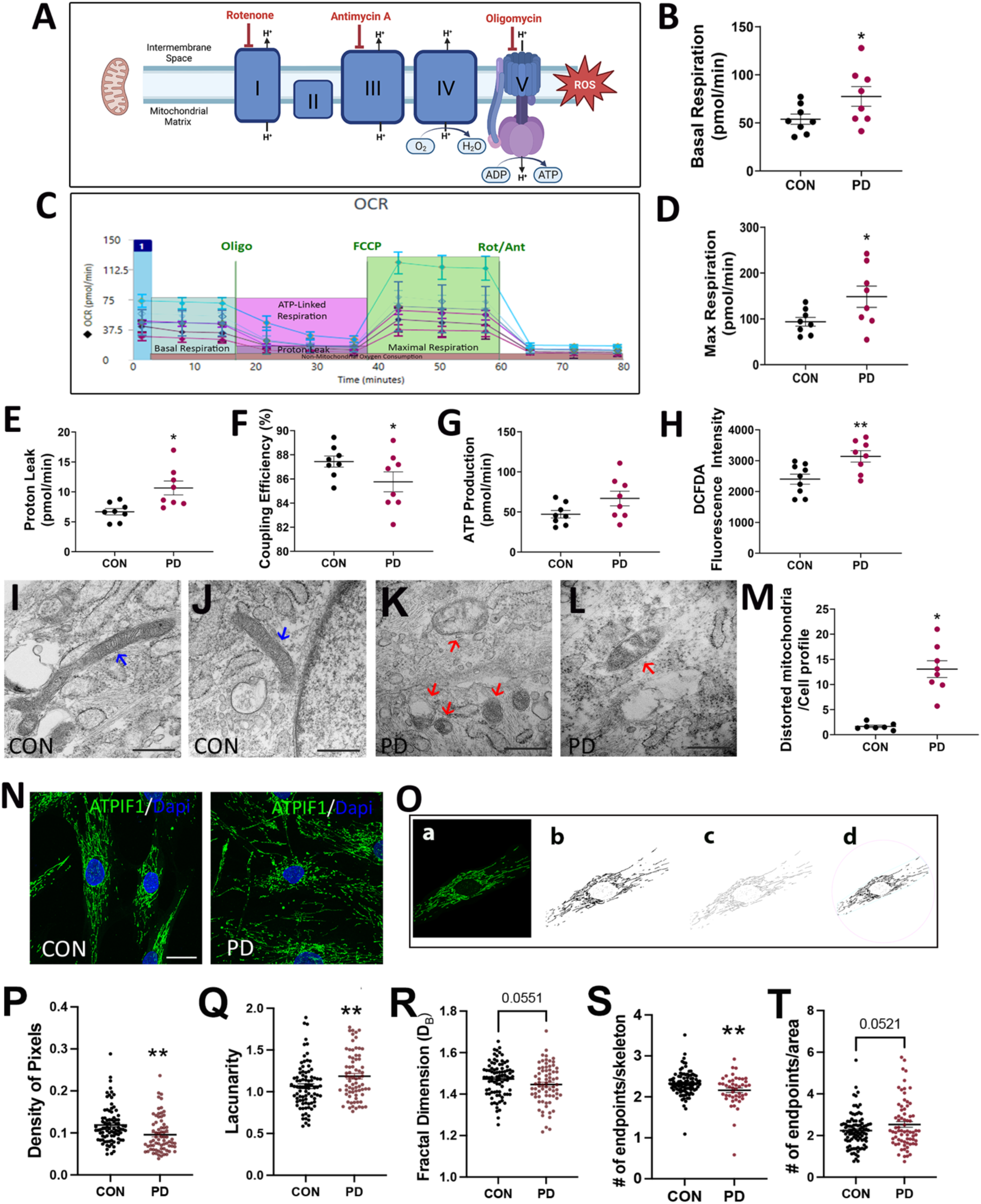
Mitochondrial-linked function analysis of the fibroblasts. (A) Schematic depiction of the mitochondrial electron transport chain and the 3 major inhibitors applied via the Seahorse assay to probe mitochondrial respiratory function. (C) shows example traces from the Seahorse Mito Stress test and some of the main parameters measured. (B-G) show results comparing the different mitochondrial function outputs obtained, particularly, basal respiration (B), maximal respiration (D), proton leak (E), coupling efficiency (F), and ATP production (G). Results from a DCFH-DA assay, showing differences in the production of gross ROS levels between CON and PD fibroblasts is in (H). TEM images showing typical mitochondria seen in CON (I, J; red arrows point to long tubular mitochondria) and PD (K, L; blue arrows point to smaller, swollen distorted mitochondria) cultures. Quantification of distorted mitochondria in CON and PD cells is shown in (M). Mitochondria were tagged via anti-ATPIF1 immunostaining (N), and an Image J pipeline was applied to threshold and skeletonize the ATPIF1 epifluorescence images allowing for characterization of mitochondrial networks (O). The major steps in the pipeline involved (a) single cell is isolated in image j, (b) image is converted to binary image, (c) binary image is skeletonized for the skeleton analysis resulting in the branch length and endpoints measurements, and (d) binary Image is run through fraclac which gives the lacunarity and pixel density. Mitochondrial properties were assessed by estimating the density of pixels (P), lacunarity (Q), fractal dimension (R), and number of endpoints (S, T). Scale Bars: (I-L) = 500nm, (N) = μm. *p<0.05, **p<0.01; Mean ± SEM, Unpaired t-tests, n = 8-9 independent lines/group. Data from the ATPIF1 mitochondrial network analysis is presented from 72-95 individual fibroblasts across 8-9 PD and CON lines to account for cell-cell variability in addition to line differences.

Using the Mito Stress test, we found that PD cells had significantly increased basal respiration and maximal respiration rates (**Fig. 2B, D**; p < 0.05; Unpaired t-test). In concert with these findings, it was found that proton leak (PL) was significantly higher (**Fig 2E**; p < 0.05; Unpaired t-test), whereas coupling efficiency (CE) was significantly lower (**Fig 2F**; p < 0.05; Unpaired t-test) suggesting that the increased PL may be the cause of increased oxygen consumption rates and lower ATP reserve in PD cells. However, significant differences were not found between ATP levels in PD vs control cells (**Fig. 2G**; p < 0.05; Unpaired t-test). In addition, non-mitochondrial respiration and spare respiratory capacity were slightly higher in the PD fibroblasts although there were no significant differences (**Supp. Fig. 1A, B**). Total reactive oxygen species (ROS) levels, measured by DCFH-DA fluorescence which represents peroxide species, was also elevated in the PD cells compared to controls (**Fig. 2H**; p < 0.05; Unpaired t-test). In essence, these data indicated altered OXPHOS in the PD fibroblasts. Additionally, in terms of glycolysis, extracellular acidification rate (ECAR) measurements indicated that basal and maximal ECAR were significantly higher in the PD fibroblasts, however the OCR/ECAR ratio was not altered (**Supp. Fig. 1C-F**, p < 0.05; Unpaired t-test) suggesting no greater reliance of the PD cells on glycolysis versus OXPHOS.

Given the mitochondrial alterations noted in PD fibroblasts, we proceeded to examine the morphology of the mitochondria in more detail via transmission electron microscopy (TEM). Here, it was observed that while control cells showed normal mitochondrial ultrastructure, with expected shape, size, and intact cristae structures (**Fig. 2I, J**, blue arrows), many mitochondria in PD fibroblasts were smaller, misshaped, and lacked typical cristae (**Fig. 2K, L**, red arrows). Quantification showed that there were higher numbers of such distorted mitochondria in PD cells compared to controls (**Fig. 2M**, p < 0.05; Unpaired t-test). Supporting these data, when mitochondrial networks were visualized using ATPIF1 targeted antibodies, PD fibroblasts showed a distinctly altered mitochondrial morphology compared to controls (**Fig. 2N**). Systematic fractal analysis of the mitochondrial structure of individual cells via NIH Image J (**Fig. 2O**) indicated that PD cells on average had significantly reduced pixel density, significantly higher lacunarity, and lower fractal dimension (**Fig. 2P-R**, p < 0.01; Unpaired t-test) indicating reduced complexity and more ‘gappiness’ of mitochondrial networks. These data together suggested less contiguous and a more fragmented mitochondrial structure in the PD cells. Additionally, skeleton analysis showed that the number of endpoints/skeleton was significantly reduced, while the number of endpoints/area was increased, in PD mitochondria, again suggesting diminished complexity and greater fragmentation (**Fig. 2S, T**, p < 0.01; Unpaired t-test). Taken together, the mitochondrial structural and functional data implied a baseline mitochondrial dysfunction in the PD fibroblasts.

### PD fibroblasts display autophagic dysregulation

Impaired autophagy is known to be an important process contributing to PD pathology, as evidenced by the collection of toxic, aggregate-prone, intracytosolic proteins in afflicted cells (Cuervo et al., 2010; Menzies et al., 2015). Our previous studies, as well as others, have shown autophagic changes in fibroblasts from both genetic and sporadic PD subjects (Galvagnion *et al*., 2022; Gonzalez-Casacuberta *et al*., 2019; Teves *et al*., 2017). From this perspective, we examined autophagy in the fibroblast lines, particularly in relation to the macroautophagy pathway (**Fig. 3A**). In particular, we examined the expression of the two standard autophagy markers, LC3 and p62, using western blotting. These experiments showed that LC3II levels were generally lower in the PD fibroblast lines as compared to controls (**Fig. 3B, D**; p > 0.05). To understand whether the trend towards lower LC3II was due to the reduced production or increased degradation of LC3II, the cells were treated with a combination of ammonium chloride and leupeptin (NH4Cl/Leup, lysosomal inhibitors) to measure autophagic flux (**Fig 3A**). Upon this treatment, an increased accumulation of LC3II was noted in PD cells (**Fig. 3B, D**) indicating higher LC3II degradation/turnover. In terms of p62, PD cell lines showed slightly higher p62 expression at baseline, compared to controls (**Fig. 3B, F**; p > 0.05). P62 expression also considerably increased upon exposure to a NH4Cl/Leup in both PD and control lines. Furthermore, under starvation conditions, greater LC3 expression was observed in both PD and control lines, which was further exaggerated by treatment with NH4Cl/Leup (**Fig. 3C, E**). P62 levels on the other hand were reduced at baseline upon starvation, in both control and PD lines although the levels were lower on average in the PD lines (**Fig. 3C, G**). Treatment with NH4Cl/Leup resulted in significant P62 increases in both control and PD lines, with greater increases seen in the control cells. All in all, these data suggested the presence of higher basal autophagy, potentially associated with a mild block, in PD fibroblasts.

**Figure 3:**
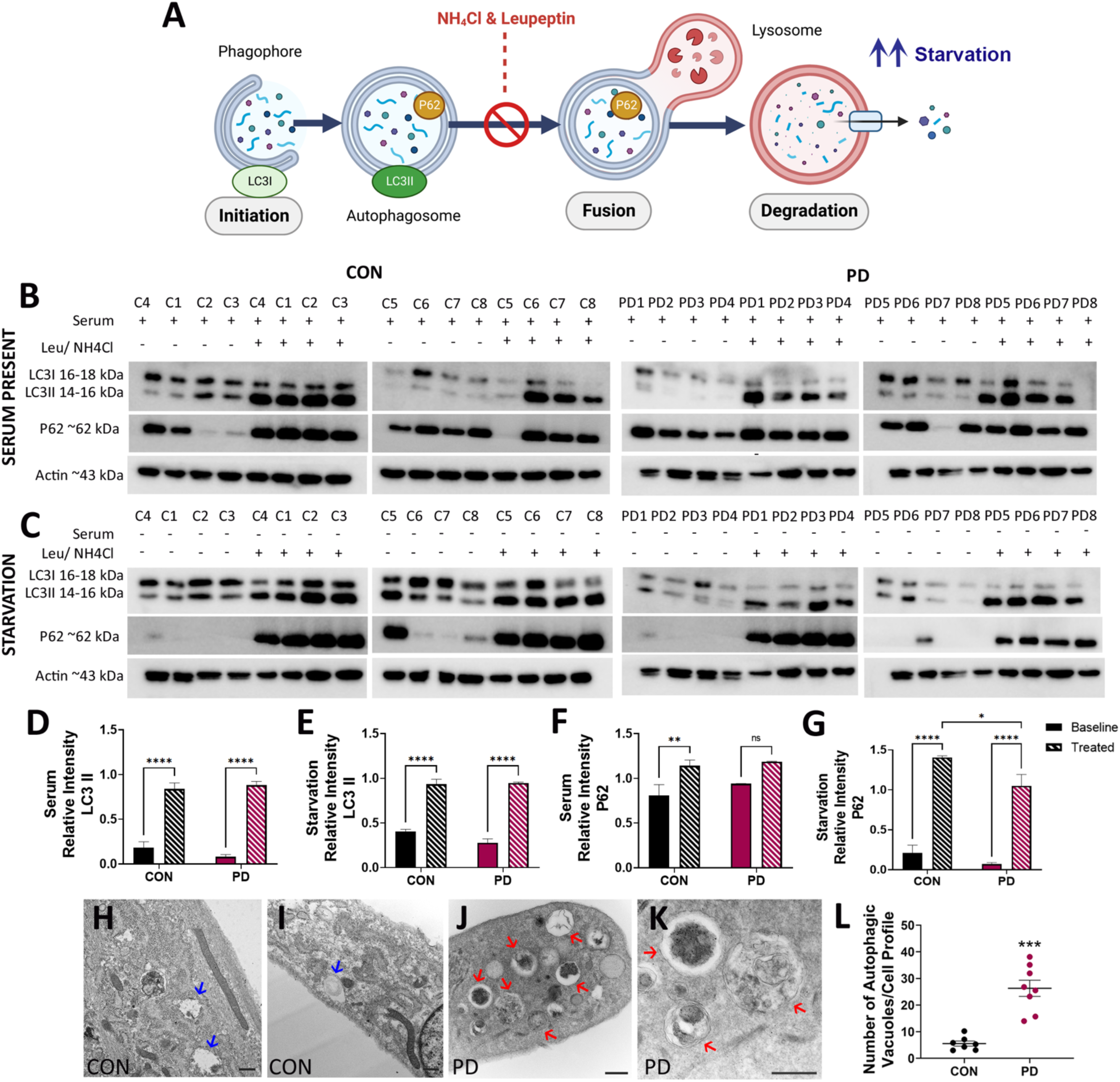
Autophagic alterations in PD fibroblasts. A schematic depicting the main phases of the macroautophagy pathway, and its classic activators and inhibitors, is in (A). Expression of the standard autophagy markers, LC3 and p62, were studied under baseline (B: serum present) and starvation (C: serum absent) conditions, with or without the lysosomal inhibitors NH4Cl/Leup, via western blotting. Densitometric quantification of the blots is shown in (D-G). (H-L) contains representative TEM images comparing the presence of autophagic structures (autophagosomes and autolysosomes) in CON (H, I; blue arrows point to autophagic vesicles) vs PD (J, K; red arrows point to several autophagic vesicles with varied cargoes) cells, and the associated quantification is in (L). Scale Bars: (H-L) = 500nm. *p<0.05, **p<0.01, ***p<0.001; Mean + SEM, Unpaired t-tests in (L), two-way ANOVAs in (D-G), n=7-8 independent lines/group.

To further probe the autophagic changes, we examined the fibroblasts in more detail via TEM. It was seen that while control cells showed the presence of some autophagic vesicles (AVs, **Fig. 3H, I**; blue arrows), sporadic PD fibroblasts exhibited a significantly increased collection of autophagic structures with varied cargo (**Fig. 3J, K**, red arrows). Quantification revealed a significant accumulation of AVs in the sporadic PD cells than in the control cells (**Fig. 3L**, p < 0.001, Unpaired t-test). These results suggested an upregulation of autophagic activity in the PD fibroblasts compared to control cells.

### Characterization of the growth, differentiation, and morphology of midbrain DA neurons generated from matched iPSCs derived from the same fibroblast lines

We next differentiated a group of matched iPSC lines (four control and four PD) generated from a subset of the fibroblast lines used in the first part of this study. Specifically we derived midbrain DA neurons, using a modified floor plate-based approach involving dual SMAD inhibition by adapting a previous method (**Fig. 4a-d; schema in Methods section**) (Chambers *et al*., 2009; Kriks *et al*., 2011). A more detailed description of the differentiation method has been included in the **supplementary methods** section. The cells were characterized for various neuronal and dopaminergic differentiation markers at 25- and 50-days post-differentiation. As shown, at day 25, robust differentiation into Tuj1^+^ neurons was observed (**Fig. 4A, C**). Quantification revealed that on average, the percentage of Tuj1^+^ cells was significantly lower in the PD cultures compared to controls (**Fig. 4E**; p < 0.05; Unpaired t-test). It was also found that about 50% of cells were expressing dopaminergic marker, Tyrosine Hydroxylase (TH), at this stage in control cultures, whereas a lower (~42%) of PD cells were TH-positive (**Fig. 4B, D, F**). We also assessed other markers indicative of midbrain DA neuron specification, mainly Lmx1a and Foxa2. As seen, although strong Lmx1a expression was seen (**Fig. 4G-N**), there were significantly lower number of Lmx1a^+^ neurons in the PD cultures vs controls (**Fig. 4O**, p < 0.05; Unpaired t-test). Similarly, while most TH cells were also Foxa2^+^, a lower percentage was found in the PD cultures (**Fig. 4P-W, X**).

**Figure 4:**
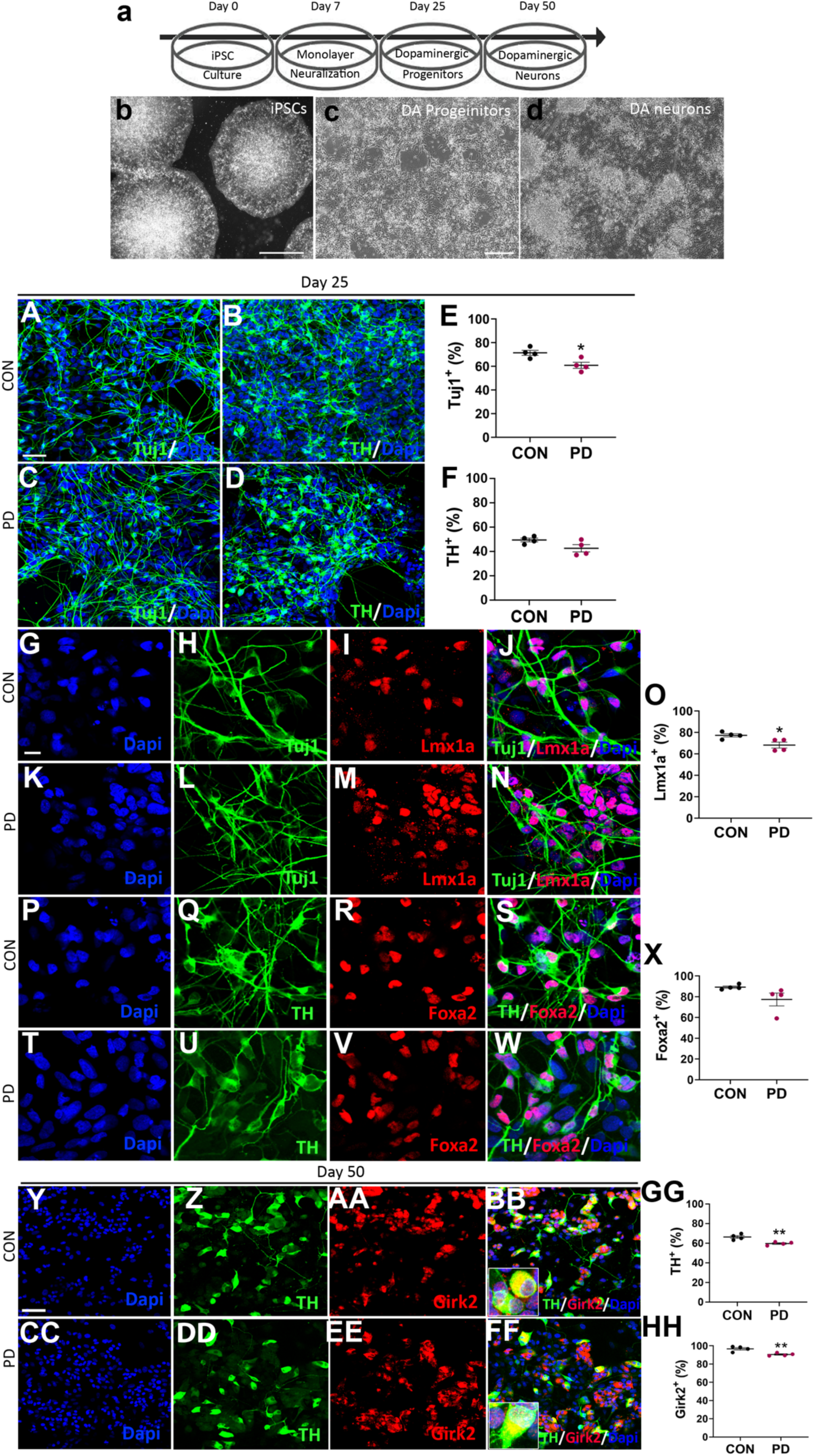
Generation and characterization of matched iPSC-DAN lines from the patient fibroblasts. Using a modified dual-SMAD inhibition protocol, iPSCs were differentiated and matured into midbrain DA neurons (a). Phase contrast images are depicted at initiation of differentiation (b – iPSC colonies), DA progenitor stage (c – Day 25) and maturing DA neurons (d – Day 50). CON and PD cells were immunostained with the neuronal marker, Tuj1 (A, C), and midbrain DA neuron marker, TH (B, D) at Day 25. Quantification of the efficiency of Tuj1^+^ and TH^+^ neuron production is shown in (E, F). Expression of the standard midbrain neuron markers Lmx1a and Foxa2 was also assessed at this stage (G-W, O, X). At day 50, TH/Girk2 co-expression was examined to determine mature A9-type midbrain DA neuron generation efficiency (Y-FF, GG, HH). High magnification views of the TH/Girk2 double-positive cells are in the insets in (BB) and (FF). Scale Bars: (b) = 1000μm, (c-d) = 100μm, (A-D, G-W, Y-FF) = 20μm. *p<0.05, **p<0.01; Mean ± SEM, Unpaired t-tests, n=4 independent lines/group.

At day 50 post-differentiation, we examined the cells for the expression of TH and Girk2. As known, TH/Girk2^+^ co-expression specifically marks mature A9 subtype DA neurons of the lateral tier of the substantia nigra (SN) that particularly degenerate in PD (Barker *et al*., 2015). It was found that while there was efficient differentiation into A9 DA neurons (**Fig. 4Y, Z, AA-FF**), there were lower percentage of TH^+^ and TH/Girk2 double-positive cells in PD vs control cultures (**Fig. 4 GG, HH**; p < 0.01, Unpaired t-tests). Compared to ~66% and 96% in control cultures, only 59% and 89% of cells were TH^+^ and TH/Girk2^+^, respectively in the PD cultures. When the cellular viability was assessed at day 50 post-differentiation (**Fig. 5A, B** shows representative phase images of PD and control neuron cultures), it was found that the PD cells on average exhibited less viability than controls although these differences were not statistically significant (**Fig. 5C**). We also examined the morphology of the cells by comparing MAP2 stained neurons in control and PD cultures (**Fig. 5D-E, G-H**). It was seen that in contrast to controls, PD neurons appeared smaller and had lesser neurite extensions. When the soma size and the number of neurites was measured (NIH Image J software), it was confirmed that the PD neurons indeed had significantly smaller soma sizes (**Fig. 5F**, p < 0.001, Unpaired t-test) as well as significantly lesser number neurites projecting from their cell bodies (**Fig. 5I**, p < 0.0001, Unpaired t-test).

**Figure 5:**
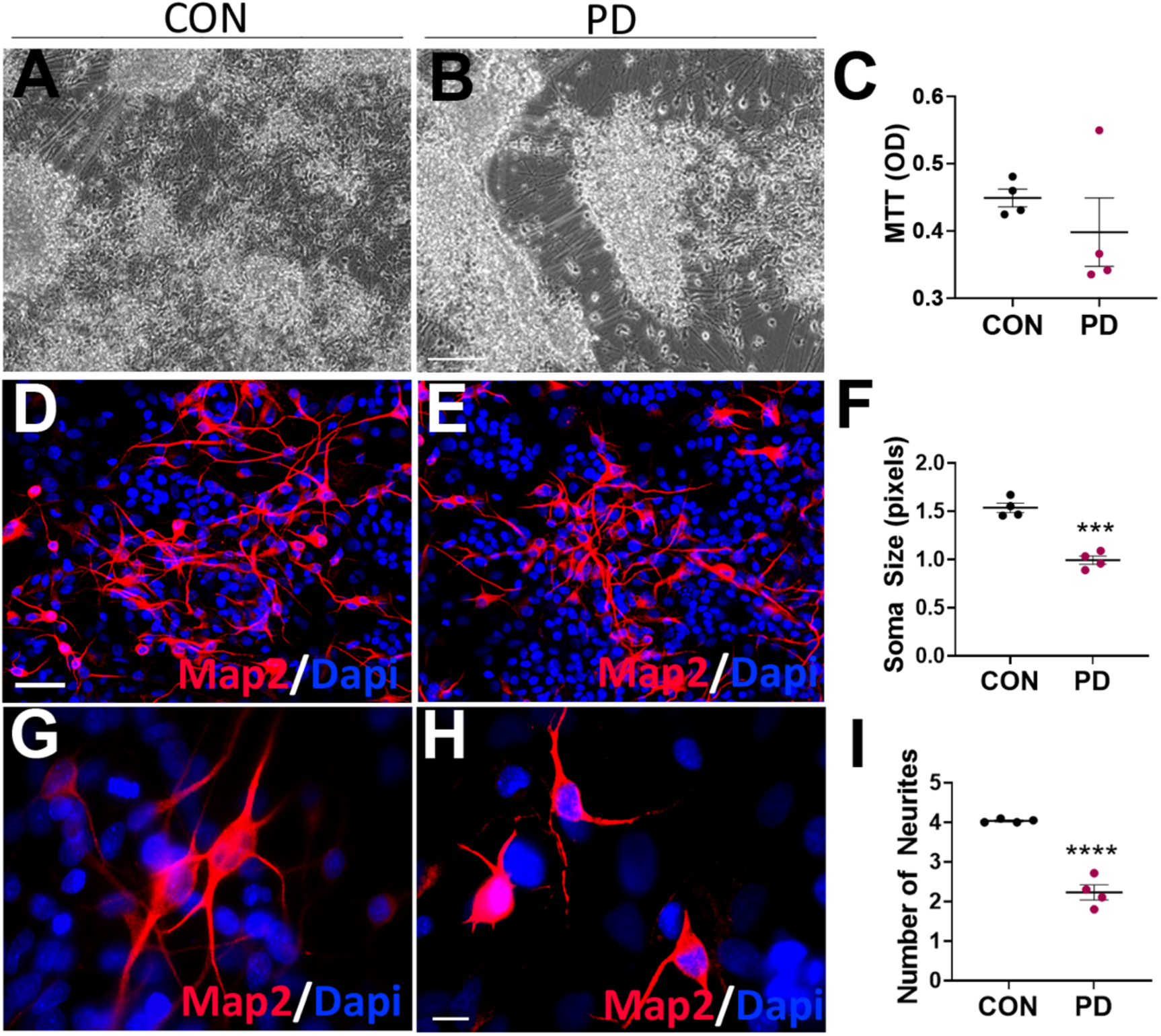
Assessment of the viability and morphological features of matched iPSC-DAN. Comparative phase images of CON and PD fibroblast-based iPSC-DAN, at Day 50 post-differentiation, is displayed in (A, B). Results from a MTT assay comparing the viability of CON vs PD iPSC-DAN is in (C). Morphological assessment was conducted on Map2 immunostained cells (D-E, G-H show higher magnification views). In particular, the soma size (F) and the number of neurites/cell (I) were estimated via Image J. Scale Bars: (A-B) = 100μm, (D-E, G-H) = 20μm. *p<0.001 ***p<0.0001; Mean ± SEM, Unpaired t-tests, n=4 independent lines/group.

### Mitochondrial structure and function are highly compromised in the PD DA neurons

Similar to the fibroblasts, the mitochondrial function of the midbrain DA neurons was examined using the Seahorse Mito Stress Test. It was found that the basal respiration of the PD DA neurons was significantly reduced compared to controls (**Fig. 6A**; p < 0.05, Unpaired t-test). Maximal respiration and PL were also notably lower in the PD DA neurons (**Fig. 6B, C**). Furthermore, there was a significant drop in ATP production in the PD DA neurons (**Fig. 6D**; p < 0.05, Unpaired t-test, with no significant differences seen in CE values (**Fig. 6E**). Non-mitochondrial respiration and spare respiratory capacity were also compromised in the PD cells vs controls (**Supp. Fig. 2A, B**; p < 0.05, Unpaired t-test). Associated with these changes in mitochondrial respiration rates, a significant elevation of total ROS levels was also found in the PD DA neurons (**Fig. 6F**; p < 0.001, Unpaired t-test). Finally, basal ECAR rates were significantly lower in the PD DA neurons, however, the basal OCR to ECAR ratio although higher for the PD cells was not statistically different from the control cells (**Supp. Fig. 2C, D**; p < 0.05, Unpaired t-test). The maximal ECAR was reduced in the PD cells with no discernable changes in the maximal OCR to ECAR ratio (**Supp. Fig.2E, F**). Overall, these data suggested a trend towards higher reliance on OXPHOS (than glycolysis) in the PD DA neurons.

**Figure 6:**
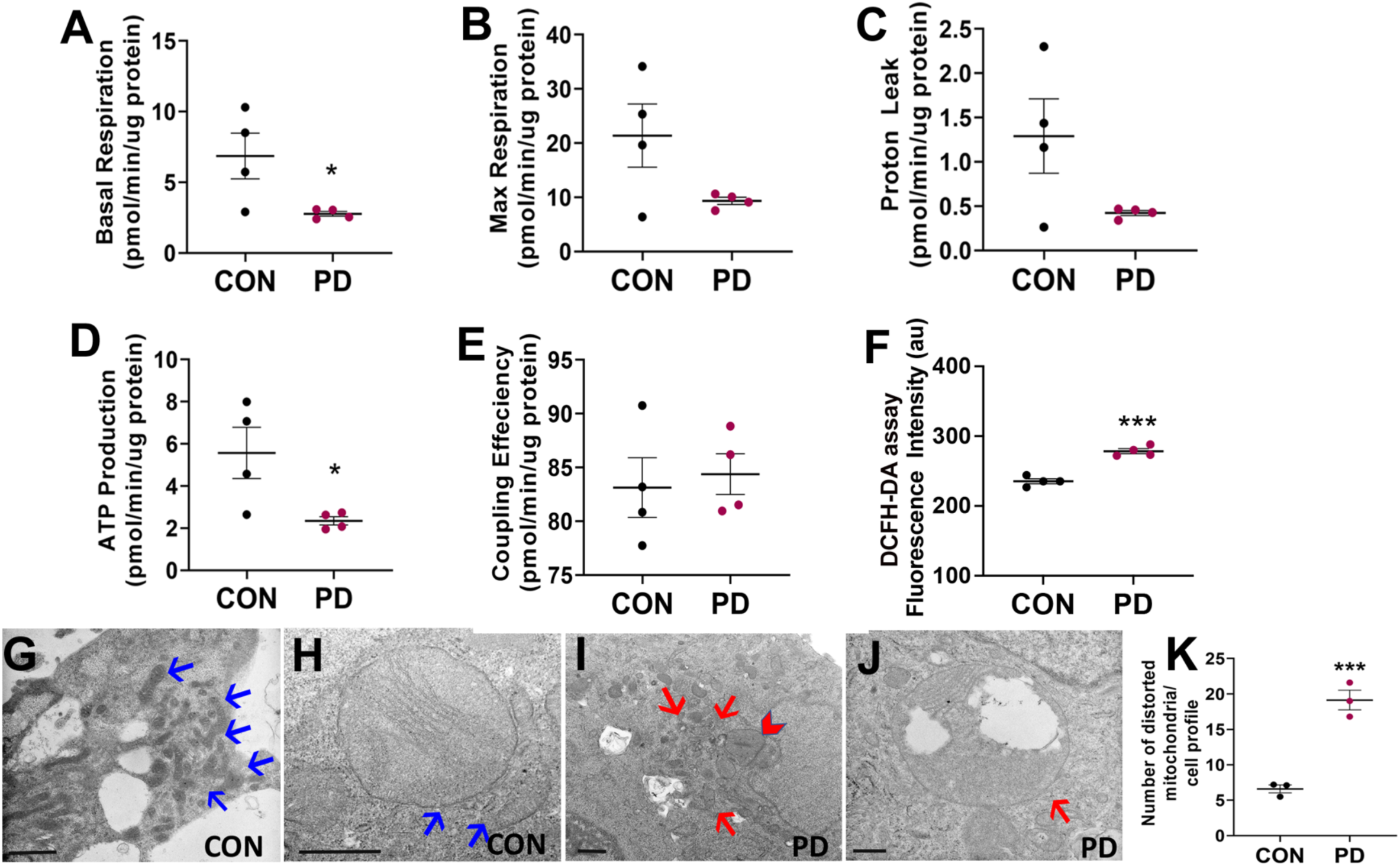
Mitochondrial dysfunction in the PD iPSC-DAN. (A-E) shows results from a Seahorse Mito Stress test, specifically basal respiration (A), maximal respiration (B), proton leak (C), ATP production (D) and coupling efficiency (E), performed on Day 50 DA CON and PD iPSC-DAN. Total cellular ROS level comparisons (DCFH-DA assay) between the CON and PD iPSC-DAN is in (F). TEM images showing the observed ultrastructure of mitochondria in CON and PD iPSC-DAN (G, H, red arrows point to normal mitochondria; I, J, red arrows point to distorted mitochondria, red arrowhead – swollen mitochondrion, vacuolating mitochondrion is seen in J). Quantification of the number of distorted mitochondria in PD vs CON iPSC-DAN is in (K). Scale Bars: (G-J): 500nm. *p<0.05, p<0.001; Mean ± SEM, Unpaired t-tests, n=4 independent lines/group.

When the subcellular morphology of the mitochondria was examined via TEM, it was observed that while control cells showed normal mitochondrial ultrastructure (**Fig. 6G, H**, blue arrows), mitochondria in the PD neurons showed altered shape, size and disruption of the typical cristae arrangement (**Fig. 6I**, red arrows, red arrowhead – swollen mitochondrion, **J** shows a vacuolating mitochondrion). Upon quantification, it was determined that there were significantly more such distorted mitochondria in PD cells compared to controls (**Fig. 6K**, p < 0.001; Unpaired t-test).

### Autophagic function and alpha synuclein expression is altered in the PD DA neurons

Autophagy, an essential catabolic mechanism for degradation of misfolded proteins and damaged organelles, is known to be affected in PD brain regions (Malkus *et al*., 2009). Thus, western blotting was conducted to investigate autophagy and the expression of alpha synuclein (aSyn), a known target of autophagy, in the DA neurons. It was seen that LC3II expression did not differ significantly at baseline in between control and PD neurons (**Fig. 7A, B**). However, treatment with the lysosomal inhibitor combination of NH4Cl/Leup resulted in a more significant increase in LC3II in PD cells than controls indicating a potentially higher LC3II degradation/turnover. In terms of p62 expression, PD cand controls did not show statistically significant differences at baseline. However, upon exposure to NH4Cl/Leup, p62 expression increased significantly in the control cells, whereas PD cells showed only a non-significant elevation (**Fig. 7A, C**). These data suggested the presence of higher basal autophagy and greater autophagic load in PD DA neurons.

**Figure 7:**
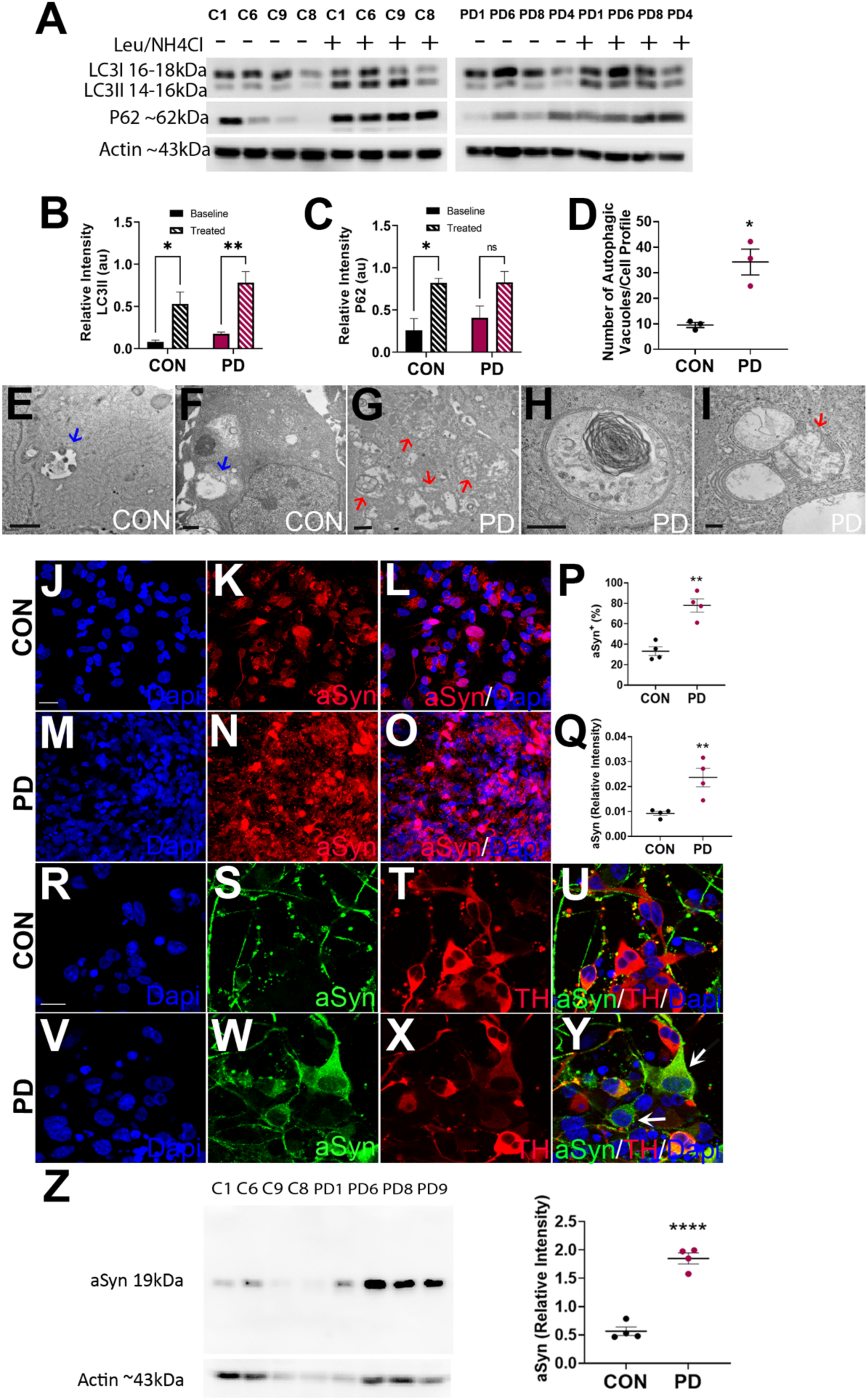
Autophagy assessments in the fibroblast matched iPSC-DAN. (A) shows western blot data from CON and PD iPSC-DAN showing LC3 and P62 turnover at baseline and after treatment with a combination of NH4Cl/Leup. Densitometric quantification is in (B, C). Representative TEM images of CON and PD iPSC-DAN, showing the relative presence of autophagic vesicles is in (E-I), with the associated quantification in (D). Red arrows point to autophagic vesicles in CON cells and blue arrows indicate those in PD cells. (H) shows a typical double membraned autophagosome in a PD iPSC-DAN, whereas (I) shows autolysosomes with different constituents, including engulfed mitochondria (red arrow in I). (J-O) and (R-Y) show comparative images of CON vs PD iPSC-DAN immunostained with antibodies against aSyn and aSyn/TH, with associated quantification in (P, Q). Expression of aSyn was also assessed via western blotting (Z). Scale Bars: (E-I) = 500nm, (J-L, M-O) = 20μm, (R-Y) = 20μm. *p<0.05, **p<0.01, ****p<0.0001; Mean ± SEM, two-way ANOVAs in (B, C), Unpaired t-tests in (D), n=4 independent lines/group.

Moreover, supporting the western blotting data, TEM analysis showed that although control cells contained some autophagic vesicles (**Fig. 7E, F**; blue arrows), sporadic PD neurons exhibited a striking accumulation of autophagic structures in their cytoplasm (**Fig. 7G**, red arrows). As shown, typical double membraned autophagosomes (**Fig. 7H**) and autolysosomes with different cargoes (**Fig. 7I**, red arrow points to engulfed mitochondria) were observed. When the number of autophagic vesicles were enumerated, as expected, significantly higher numbers were seen in PD neurons (**Fig. 7D**, p < 0.05, Unpaired t-test). Essentially, all these data suggested dysregulated autophagy in PD DA neurons.

In terms of the PD-relevant protein, aSyn, immunocytochemical analysis revealed that PD cultures had more aSyn than control cultures (**Fig. 7J-O**). In fact, both the number of aSyn expressing cells and the relative intensity of aSyn expression was higher in the PD cultures than in controls (**Fig. 7P, Q**; p < 0.01, Unpaired t-test). In addition, it was observed that TH^+^ DA neurons in PD cultures had higher aSyn expression within their cell bodies compared to controls (**Fig. 7R-Y**). Interestingly, many PD neurons with high intracytoplasmic aSyn, showed reduced TH immunoreactivity suggesting dopaminergic dysfunction (white arrows in **7Y**). Furthermore, increased aSyn expression in the PD DA neurons was supported by data from western blotting studies, which showed a similar trend (**Fig. 7Z**). Given the autophagic dysregulation seen in PD neurons, the buildup of aSyn in these cells may be attributed to inefficient autophagic clearance [34]. Given these data, as a comparison, we also assessed aSyn expression in the fibroblasts. Immunocytochemistry revealed higher aSyn in the PD fibroblasts than controls, although the levels were seen to be generally low (**Supp. Fig. 3A-C**). Nonetheless, western blotting and dot blot analysis did not detect aSyn in any of the lines.

### PD DA neurons express different electrophysiological properties from controls

The electrophysiological activity of PD and control neurons was analyzed at rest and after current injections to attempt to induce spiking. The different types of spiking activity profiles seen are depicted in **Figure 8A**. Overall, it was found that 77% of PD neurons were inactive, showing no spiking either spontaneously at rest or when induced with current steps. In contrast, only 65% of control neurons were inactive, suggesting that the PD neurons were less active as a population (**Fig. 8B**). Although similar proportions of control and PD neurons had spontaneous spiking (16% vs. 15%), PD neurons had a much smaller proportion of cells with inducible spiking (19% vs. 6%). All inducible neurons but one showed spiking after a −30pA current step while one showed spiking after a +30pA step (**Fig. 8A**, bottom). This suggests the Na^+^ channels were in an inactive state at baseline. Furthermore, both groups had fairly depolarized resting membrane potentials for mature neurons. However, control neurons had significantly more negative resting membrane potentials than PD neurons (**Fig. 8C**, control: −29 ± 2.73; PD: −22 ± 1.98 mV; p<0.05, one-way ANOVA with SNK posthoc test). There were no differences in resting membrane potential between the four cell lines of either control or PD. In addition, mean resting membrane potentials of spontaneously active cells were significantly lower than those of inactive cells (spontaneous: −40 ± 3.25; non-active: −20 ± 1.59 mV, p<0.001; two-way ANOVA with SNK posthoc test). When broken down by activity and group, spontaneously active neurons had lower resting membrane potentials than inactive neurons in control (**Fig. 8D**, spontaneous: −39 ± 5.3; non-active: −24 ± 2.65 mV, p=0.012) and PD cultures (spontaneous: −42 ± 4.9; non-active: −18 ± 2.1 mV, p<0.001; two-way ANOVA with SNK posthoc test).

**Figure 8:**
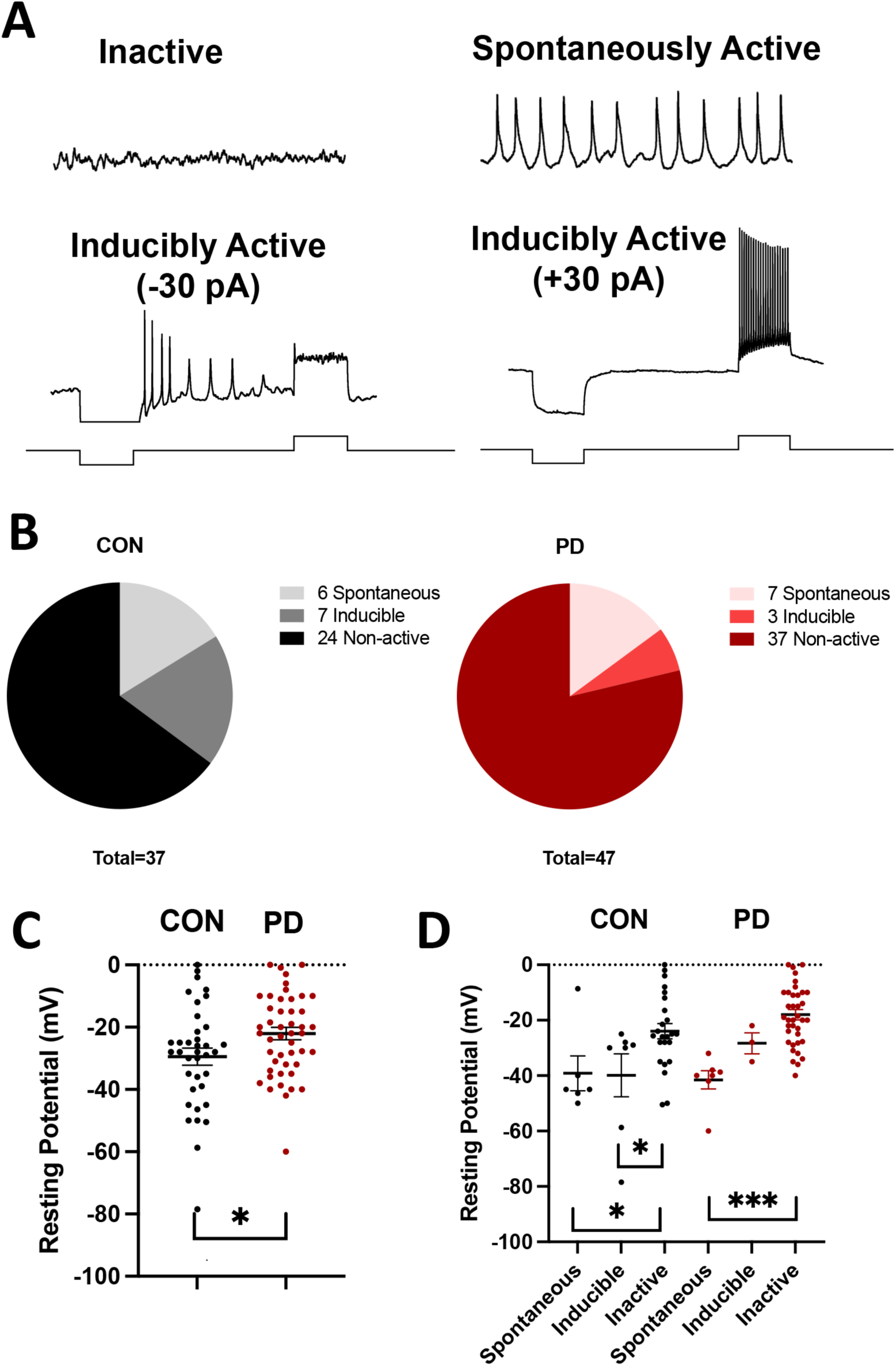
Electrophysiological analysis of CON and PD iPSC-DAN. Day 55 iPSC-DAN were subjected to whole cell recordings and analyzed for spontaneous and evoked action potentials. Representative current-clamp recordings of inactive, spontaneously active, and inducible active neurons (with −30 pA current step, and +30 pA step) are in (A). Summary of type of cell activity between CON and PD neurons is shown in (B). (C) depicts a comparison of the mean resting potential between CON and PD neurons. Dots represent data values for individual cells. (D) displays the differences in mean resting potential by activity of spontaneous, inducible, and inactive neurons between CON and PD groups. *p<0.05, **p<0.01, ***p<0.001; Mean ± SEM, one-way ANOVA with Student-Newman-Keuls posthoc test in (C), two-way ANOVA with Student-Newman-Keuls posthoc test in (D), n=3 independent lines/group.

Since our data indicated a reduced soma size in PD neurons, we also assessed cellular capacitance. However, an analysis of cell capacitance showed no significant difference between control and PD neurons (**Table 2**: p=0.617; *see at the end of this document*). There was also no significant difference in cell capacitance between individual cell lines. Since neurons that were successfully recorded from had less variability in size as compared to the population of plated cells, cell capacitance measures may not be a good representation of the neuron sizes as a whole population.

**Table 2.** Electrophysiological data.

| Group | Parameter | Mean | 95% CI | N Value | P Value |
| --- | --- | --- | --- | --- | --- |
| Control | Capacitance | 12.05 (pF)<br>Median | -1.47 - 2.61 | 36 | 0.579 |
| PD |  | 11.84 (pF)<br>Median |  | 46 |  |
| Control | Capacitance<br>(Spontaneous) | 9.34 (pF) | -10.02 - 2.26 | 6 | 0.186 |
| PD |  | 12.99 (pF) |  | 7 |  |
| Control | Frequency<br>(Spontaneous) | 0.7 (hz) | -221.86 –<br>139.19 | 3 | 0.605 |
| PD |  | 1.13 (hz) |  | 6 |  |
| Control | Peak Amplitude<br>(Spontaneous) | 38.52 (mV) | -13.11 – 13.78 | 3 | 0.955 |
| PD |  | 38.19 (mV) |  | 6 |  |
| Control | Decay Tau<br>(Spontaneous) | 338.21<br>(ms) | -181.80 –<br>560.12 | 3 | 0.267 |
| PD |  | 149.047<br>(ms) |  | 6 |  |
| Control | Inter-event<br>Interval<br>(Spontaneous) | 1262.76<br>(ms)<br>Median | -8249.58 –<br>5349.05 | 3 | 0.629 |
| PD |  | 930.08<br>(ms)<br>Median |  | 6 |  |
| Control | Time to Peak<br>(Spontaneous) | 174.51<br>(ms) | -10.46 –<br>191.41 | 3 | 0.072 |
| PD |  | 84.04 (ms) |  | 6 |  |

Action potentials of spontaneously firing neurons were also analyzed. It should be noted that there was a large amount of variability in spiking between spontaneously active neurons. Some neurons spiked as many as 324 times during a 2-minute recording and one neuron only spiked twice. There were no differences in peak amplitude, decay tau, inter-event interval, or time to peak of spontaneous spikes between control and PD neurons (**Table 2**, *at the end of this document*).

### Cross correlation analyses of fibroblast, DA neuron, and clinical data

We compared the cellular data obtained from the fibroblasts and neurons to each other, and also correlated the cellular data from each of the cell types to some available clinical data from individual PD subjects. The scatterplots in **Figure 9**, display the relationship between the groups (control and PD) and sex for different fibroblast and DA neuron assays, side by side, to allow for pairwise comparison. The data is expressed as fold changes normalized to mean of controls, and the line in each plot represents the change between the mean value of each group. The slope of the line signifies the change between control and PD groups. The shaded area represents a CI for the placement of the line, which is directly linked to the dispersion in the data points.

**Figure 9:**
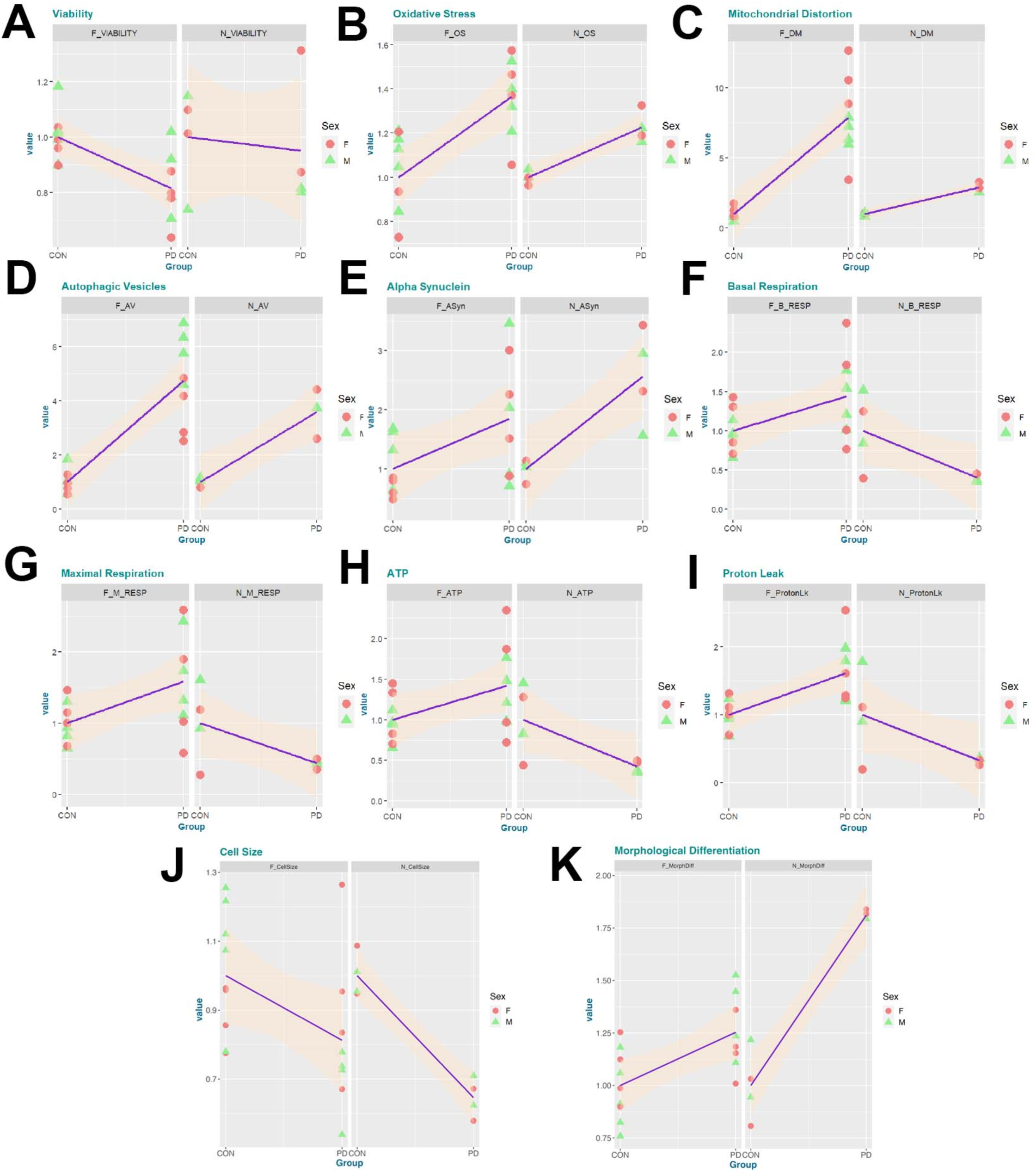
The scatterplots display a pairwise comparison of the relationship between CON and PD groups and sex for fibroblast and neuron cellular assays. Specifically, results shown compare fibroblast to neuron viability (A, MTT assay), oxidative stress (B, DCFH-DA assay), number of distorted mitochondria (C, electron microscopy), number of autophagic vesicles (D, electron microscopy), alpha-synuclein (E), basal respiration (F, seahorse assay), maximal respiration (G, seahorse assay), ATP production (H, seahorse assay), and proton leak (I, seahorse assay), cell size (J) and morphological differentiation (K, comparison of form factor in fibroblasts to the and number of neurites in neurons, parameters that measure process ramification). The data has been modified to represent fold changes compared to the mean of controls. The line in each plot represents the change between the mean value of each group, and the slope of the line signifies the change between CON and PD groups. The shaded area represents a confidence interval (CI) for the placement of the line.

As seen, the PD groups showed reduced viability and increased oxidative stress in both the fibroblasts and neurons; however, the fibroblast’s change is higher between the groups compared to neurons represented by the increased slope in the line (**Fig. 9A, B**). Similarly, the changes in number of distorted mitochondria and number of AVs trended in the same direction for both fibroblasts and neurons (**Fig. 9C, D**). While the amount of change was similar with respect to AV number, the change was higher in fibroblasts with respect to distorted mitochondrial number. Furthermore, the relative expression of aSyn was higher in both PD neurons and PD fibroblasts compared to their control counterparts, however, the slope was higher in the neurons (**Fig. 9E**). Interestingly, with respect to the mitochondrial function measures of basal respiration, maximal respiration, PL and ATP production, the directionality of changes was contrary between the fibroblasts and DA neurons (**Fig. 9F-I**). For example, PD neurons had lower basal respiration, maximal respiration, PL and ATP production, compared to control neurons (negative slopes), whereas fibroblasts showed the opposite trend (positive slopes). The amount of change in the positive or negative slopes were similar. Lastly, the cell size (**Fig. 9J**) and morphological differentiation (**Fig. 9K**). was reduced in both PD fibroblasts and the DA neurons.

Correlation matrices (Heat Maps) were generated between specific clinical and cellular data for the fibroblasts (**Fig. 10A**, Fibroblast-Clinical) and neurons (**Fig. 10B**, Neurons-Clinical). The third correlation matrix is between the particular cellular assays of neurons and fibroblasts (**Fig. 10C**, Fibroblast-Neuron). All these plots show negative correlations in dark red and positive correlations in dark grey. Correlation values vary between the range of negative one and positive one, [-1,1], and the matrix depicts correlations between all possible pairs of variables. The diagonal shows the correlation of each variable with itself, which is a perfect 1. When two variables have no relation to one another they would show zero correlation, depicted with the color white in all three plots. Therefore, colors that are faintly visible only show a slight or negligible correlation and are close to zero. In the fibroblast-clinical correlations heat map (**Fig. 10A**), interesting positive associations between numbers of AVs and UPDRS scores, as well as mitochondrial function (basal and max respiration) and UPDRS scores, can be seen. Also, positive correlations between oxidative stress and CSF aSyn, CSF Abeta, and age are seen. In contrast oxidative stress is negatively correlated to UPDRS3_ON scores. With respect to the neuron-clinical plot (**Fig. 10B**), positive correlations between number of AVs and disease duration, age, CSF aSyn, CSF Abeta and neuronal aSyn are observed. Positive correlations are seen between neuronal aSyn and CSF aSyn and CSF Abeta. On the other hand, both the efficiency of DA neuron generation and neuronal maximal respiration are negatively correlated with number of AVs, CSF aSyn, CSF Abeta and neuronal aSyn. In terms of the fibroblast-neuron correlation matrix (**Fig. 10C**), as expected, positive correlations between data from similar fibroblast and neurons assays are seen, except for the mitochondrial function data where the correlations are negative. In terms of correlations of data within each cell type, the population doubling time of fibroblasts is negatively correlated with most other fibroblast measures. The efficiency of DA neuron generation is positively correlated with neuronal basal and maximal respiration levels and negatively correlated with the number of AVs and aSyn expression. Surprisingly, a strong negative correlation between age and neuronal viability was seen, but there was no notable relationship between age and viability in the fibroblasts (**Fig. 10A, B; Supp. Fig. 4A, B**). Additionally, disease duration (DD) was negatively correlated with viability in both the fibroblasts and the neurons (**Fig 10A, B; Supp. Fig. 4C, D**).

**Figure 10:**
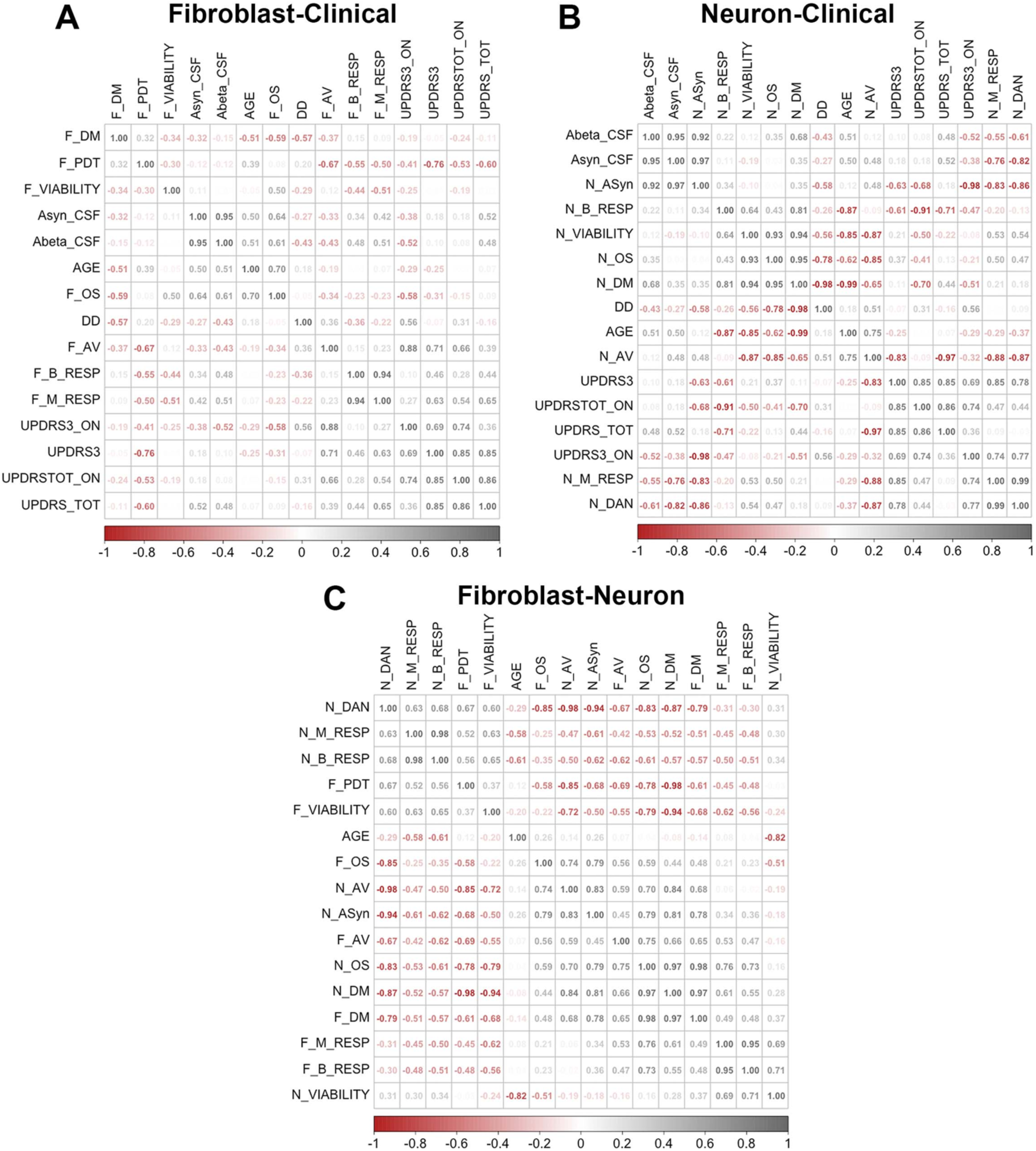
Correlation matrices comparing clinical data obtained at the time of sample collection to results from the fibroblast assays (A, Fibroblast-Clinical), and the neuronal assays (B, Neuron-Clinical) are shown. (C, Fibroblast-Neuron) displays the correlation matrix between cellular data from neurons and cellular data from the fibroblasts. These plots show negative correlations in dark red and positive correlation in dark grey. Age (age at sample collection) and DD (disease duration) are included in all the correlation matrices. The matrices depict the correlation between all possible pairs of variables. The diagonal shows the correlation of each variable with itself which is a perfect 1. Off the diagonal elements each correspond to the correlation between a pair of variables which are indicated in the axes of the plot. For this reason, the two sides of the diagonal are mirror images of each other as there are two cells in each matrix showing the same correlation between a pair of variables. The higher the correlation the darker the color. Zero correlation is shown with the color white in all three plots, therefore colors that are faintly visible only show a slight or negligible correlation and are close to zero. This is shown by a color legend at the bottom of each plot. Co-efficient were calculated based on Pearson’s correlation via Corrplot package in R.

## DISCUSSION

In essence, we report parallel neurodegenerative phenotypes in sporadic PD fibroblasts and matched iPSC-based DA neurons, *derived from the same patient*, which capture several fundamental PD mechanisms. Specifically, using this approach, we identified a range of pathological alterations in ‘aged’ PD patient fibroblasts and ‘young’ iPSC-derived DA neurons reprogrammed from the fibroblasts, that reflect disease processes relevant to both early neural development and ageing. Specific correlations between the fibroblast and neuronal features were seen which highlighted cell type- and tissue type-specific differences at play in PD. To our knowledge, this is the first time a bicameral patient-specific system consisting of both peripheral (skin fibroblast) and central/neural (iPSC-DAN) cells has been used to conduct a detailed mechanistic modeling of idiopathic PD.

The study has a few important implications. Firstly, the data indicate that basic mechanisms of PD can be expressed beyond the boundaries of the central nervous system, in peripheral cells such as skin fibroblasts – thus supporting the view of PD as a generalized metabolic disorder rather than a neuron-centric condition. Secondly, the corresponding changes in the DA neurons, derived from the fibroblasts, suggests that dysfunction in peripheral tissues would also be associated with abnormalities in the brains of respective sporadic PD patients. Thirdly, and in a broader sense, the results imply that by taking advantage of the reprogramming effect, different patient-derived cells, along the fibroblast-iPSC-neural progenitor-neuron-glia differentiation continuum, can be used as powerful models to identify cell type-specific neurodegenerative mechanisms active ‘earlier’ in the disease. By extension, utilization of these varied cell types can help build robust biomarker and drug testing platforms relevant to developing much needed early therapeutic interventions for PD.

In terms of the cellular phenotypes seen, in most cases, we found positive associations between data from similar fibroblast and neuronal assays. For instance, compared to their control counterparts, PD fibroblasts and PD iPSC-DAN showed reduced viability (MTT test), increased ROS levels (DCFH-DA test), abnormal mitochondrial structure (TEM) and function (Seahorse Mito Stress test), dysregulated autophagy (TEM and western blotting) and altered morphology (cell size and shape measurements). However, there were also notable differences. Interestingly, the pathological changes appeared to be more severe in the diseased fibroblasts than the iPSC-DAN, with greater differences seen between control and PD within this cell type. For instance, a greater reduction in viability and larger increases in oxidative stress, number of distorted mitochondria and autophagic vesicles were seen in PD fibroblasts vs PD iPSC-DAN when compared to their relevant controls. These changes could be attributed to the contribution of age-related variations in the fibroblasts; an aspect lost in the iPSC-DAN (Ivanov *et al*., 2016; Mertens *et al*., 2018). However, with respect to mitochondrial function, PD iPSC-DAN exhibited a worse phenotype than the fibroblasts. In particular, the PD iPSC-DAN had significantly lower basal and maximal respiration rates, proton leak, ATP production, and spare respiratory capacity compared to controls. The fibroblasts on the other hand showed increased basal and maximal respiration rates, proton leak, and spare respiratory capacity, and no significant changes in ATP production, compared to their controls. This occurred despite the elevations in the number of distorted mitochondria being higher in the fibroblasts compared to the neurons, suggesting that fibroblasts were able to compensate to some extent whereas the neurons were more vulnerable to the mitochondrial alterations. Also, although more distorted mitochondria were present in PD fibroblasts, the severity of the distortions (mitochondrial swelling, disruption of cristae, vacuolization, and mitophagy) were greater in PD iPSC-DAN than PD fibroblasts, supporting greater neuronal susceptibility to mitochondrial dysfunction. These data align with other studies that indicate that certain attributes of A9 midbrain DA neurons render them highly vulnerable to mitochondrial dysfunction (Greenamyre and Hastings, 2004; Matsuda *et al*., 2009; Sulzer, 2007; Surmeier *et al*., 2011).

It was also noted in the fibroblasts, that while there was increased proton leak and reduced coupling efficiency, the ATP production was not significantly different between control and PD groups. These data suggest that fibroblast PD mitochondria may have a higher capacity to generate the protonmotive force (PMF), which is the driving force for both OXPHOS and proton leak (Berry *et al*., 2018). High PMF is good for generating a robust ATP free energy, but it may also promote superoxide production. In support of this idea, higher ROS levels were seen in the PD fibroblasts compared to control cells. Also, by dissipating the PMF, the proton leak may protect against high free radical production. Thus, by increasing the proton leak PD fibroblasts maybe “uncoupling to survive.” In addition to these aspects, basal and maximal ECAR were significantly higher in the PD fibroblasts, in comparison to the iPSC PD-DAN, which showed opposite trends, indirectly pointing to different rates of glycolysis in the two cell types (although the OCR/ECAR ratios were not significantly different). All in all, these results support the notion that the fibroblasts are more able to compensate for mitochondrial impairments vs. the DA neurons. Nonetheless, additional studies will be needed to gain a fuller understanding of the dynamics of these mitochondrial processes.

The PD DA neurons also exhibited increased aSyn accumulation and altered electrophysiological profiles. Specifically, both immunocytochemical and western blotting data showed that the PD iPSC-DAN generally expressed more aSyn than control neurons. Since basal autophagy was increased in the PD iPSC-DAN, and aSyn is a known autophagy cargo, these data imply that the PD neurons maybe engaging in more autophagic activity to plausibly cope with a higher load of aSyn. On the other hand, it is known that an increased aSyn protein burden may impair autophagy, via mechanisms such as Rab1a inhibition and blockade of the high-mobility group box 1 protein, thus generating a bidirectional positive feedback loop (Huang *et al*., 2017; Winslow *et al*., 2010). However, whether autophagy dysfunction alone suffices to increase aSyn or whether aSyn is the pathogenic driver, is an important question that needs to be addressed.

Besides autophagy, it is recognized that increased aSyn can also negatively affect mitochondrial function and cytoskeletal structure. In fact, it has been reported that aSyn can localize to mitochondria and induce severe ultrastructural deformation of the inner mitochondrial cristae membranes, massive swelling and a loss of respiratory function (Ganjam *et al*., 2019). There is also evidence that aSyn interacts with actin and tubulin, as well as their specific binding proteins, to impact cytoskeletal integrity (Carnwath *et al*., 2018; Oliveira da Silva and Liz, 2020). Cytoskeletal destabilization has been implicated as a major player that paves the way for neurodegeneration in PD, and our data on cell shape and size alterations in the PD neurons (as well as fibroblasts) suggest that such a mechanism could be at play in the human PD lines (McMurray, 2000; Pellegrini *et al*., 2017).

With respect to neuronal activity, a larger proportion of PD DA neurons were found to be inactive and did not show any action potentials, either spontaneously at rest or when induced with current steps, compared to controls. Moreover, although similar proportions of control and PD neurons were spontaneously active, a much smaller fraction of PD neurons could be induced to spike. Overall, these results implied that the PD iPSC-DAN were less electrically active as a population. Interestingly, it was also found that the control neurons had significantly more negative resting membrane potentials compared to the PD neurons, providing one reason why the control neurons were more readily activated. Neurons critically depend on mitochondrial function to establish membrane excitability and to execute the complex processes of neurotransmission and plasticity (Mattson *et al*., 2008). From this perspective, it is possible that the impaired mitochondrial function/ATP production observed in the PD iPSC-DAN maybe contributing to their altered electrophysiological profiles. Likewise, basic disruptions in Na^+^ and K^+^ channel dynamics could also be driving the differential activity of the control and PD neurons. Nevertheless, these hypotheses need to be carefully studied to understand the precise underlying biology.

The statistical correlational data comparing the fibroblast and neuronal outputs to different clinical measures from the patients, showed several interesting associations. Of note was the inverse relationship between age and viability in the DA neurons but not the fibroblasts. Given the greater maintenance of aging marks in the fibroblasts compared to the iPSC-DAN this was an unexpected finding. However, disease duration was negatively correlated with age with respect to both cell types, suggesting that, contributions of age, disease duration, and cell-type specificity (neuron vs fibroblast), may be important in determining the cellular phenotype. A more comprehensive study with larger sample sizes would be needed to obtain a clearer picture. Nevertheless, overall, although the data sets are relatively small, the presented statistical analyses provide an appreciation of how such methods could support a robust stratification of the patient lines with respect to both mechanisms as well as clinical measures.

In conclusion, our identification of parallel phenotypes in sporadic PD fibroblast and iPSC-based DA neurons from the same patient, creates a novel system to study both early neural and age-related peripheral influences driving the progression of PD. By lending itself to such mechanistic studies, this system also provides a strong platform for the discovery of biomarkers for the early diagnosis and stratification of PD patients. Specifically, the study confirms the utility of patient skin fibroblasts as an accessible peripheral cell system reflective of disease status through the assay of multiple distinct biologic features. The study also reveals that iPSC-DAN reprogrammed and differentiated from the fibroblasts, can spontaneously express synchronous phenotypes, providing ‘early neural correlates’ of the fibroblast changes. Accordingly, we propose that such a multi-level modeling approach, involving different peripheral as well as central iPSC-derived neural cells, can enrich perspectives obtained from other important cellular systems like directly reprogrammed cells, to support a deeper understanding of disease mechanisms and accelerate the development of rational therapeutics for early intervention in PD.

## ACKNOWLEDGEMENTS

We thank Dr Helena Morrison and Dr Wayne Willis for their expert input on the mitochondrial ATPIF1 and seahorse data. We also acknowledge Drs Tony Day and Paula Tonino for their assistance with the transmission electron microscopy work, which was performed in the Imaging Cores-Electron at the University of Arizona Research, Innovation, and Impact Core Facilities. Our thanks also to Dr Maria Sans-Fuentes, for her assistance with the statistical analysis. The cell lines used in this study were partially obtained from the Parkinson’s Progression Markers Initiative (PPMI) [www.ppmi-info.org]. PPMI, a public private partnership, is funded by the Michael J. Fox Foundation for Parkinson’s Research and corporate sponsors [https://www.ppmi-info.org/about ppmi/who-we-are/study-sponsors]. The study was supported by a Michael J Fox Foundation Grant (MJFF 18366) and UA intramural funds to LM, and National Eye Institute Grant R01-EY-026027 and NSF-1552184 to EE.

## AUTHOR CONTRIBUTIONS

MJC – Conception and design, Collection and assembly of data, Data analysis and Interpretation, Manuscript writing

AMJ – Collection and assembly of data, Data analysis and Interpretation, Manuscript writing

EC – Collection and assembly of data, Data analysis.

TM – Collection, assembly and analysis of electrophysiology data, Manuscript writing

EE – Conception and design of electrophysiology experiments, Data analysis and resources support, Manuscript editing

KB – Collection, assembly, data analysis of ATPIF1 data, Manuscript editing

ML – Experimental design, execution, and data analysis support for Seahorse experiments

LJM – Experimental design, data interpretation and resources support for Seahorse experiments, Manuscript editing

SP – Statistical data analysis and interpretation, Manuscript editing

DB – Statistical data analysis and interpretation, Manuscript editing

LM – Overall conception and design, Collection and assembly of data, Data analysis and Interpretation, Manuscript writing, Financial support, Final approval of manuscript.

## SUPPLEMENTARY FIGURES

**Supplementary Figure 1:**
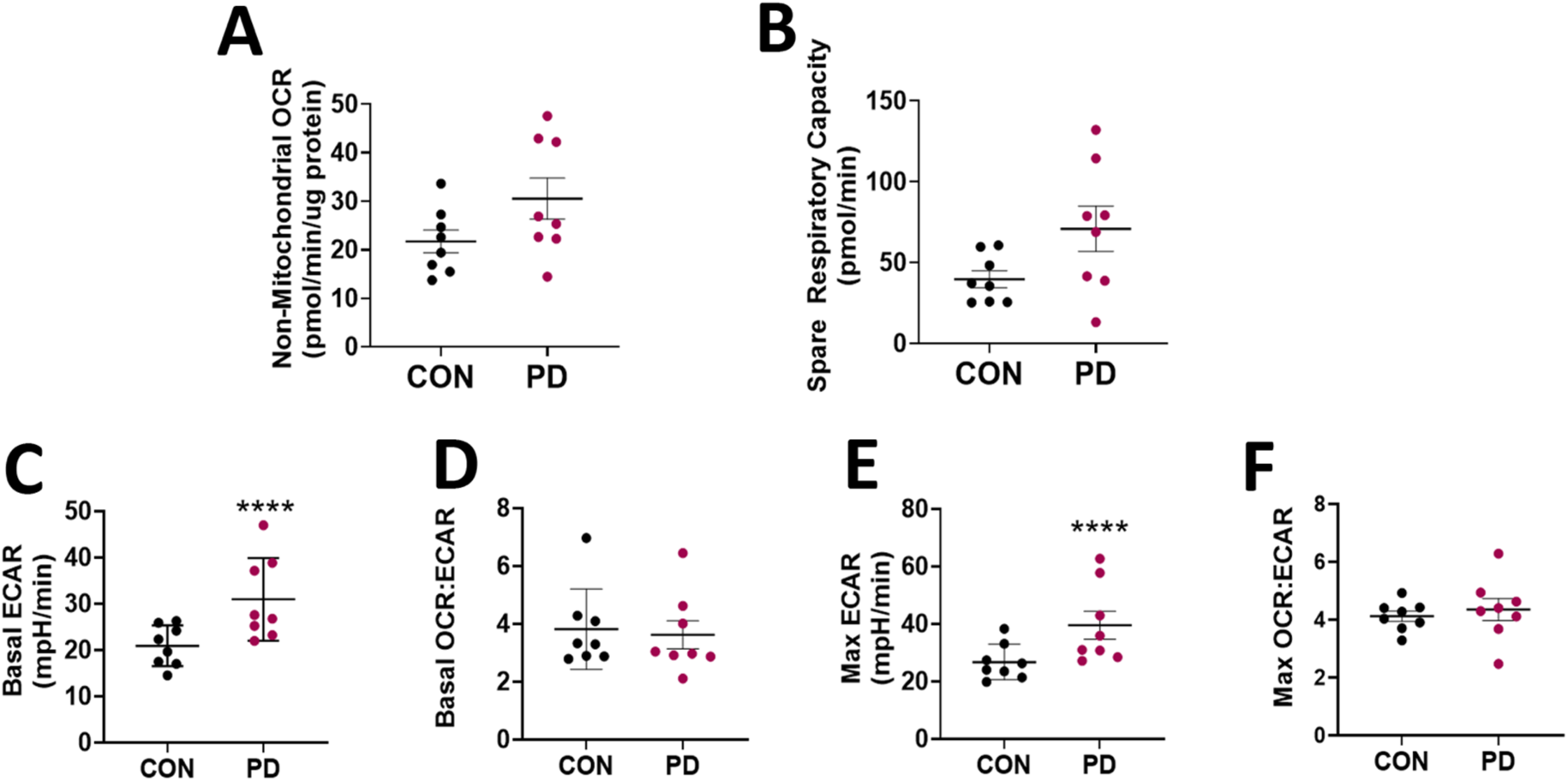
Seahorse Mito Stress analysis of fibroblasts. Additional OCR and ECAR measurements were obtained via the Seahorse Mito Stress test. Non-mitochondrial linked oxygen consumption (A) and spare respiratory capacity (B) were compared between CON and PD cells. Basal ECAR and Maximal ECAR (C, D), and OCR/ECAR ratios were also calculated (E, F). ****p<0.0001; Mean ± SEM, Unpaired t-tests, n=8 independent lines/group.

**Supplementary Figure 2:**
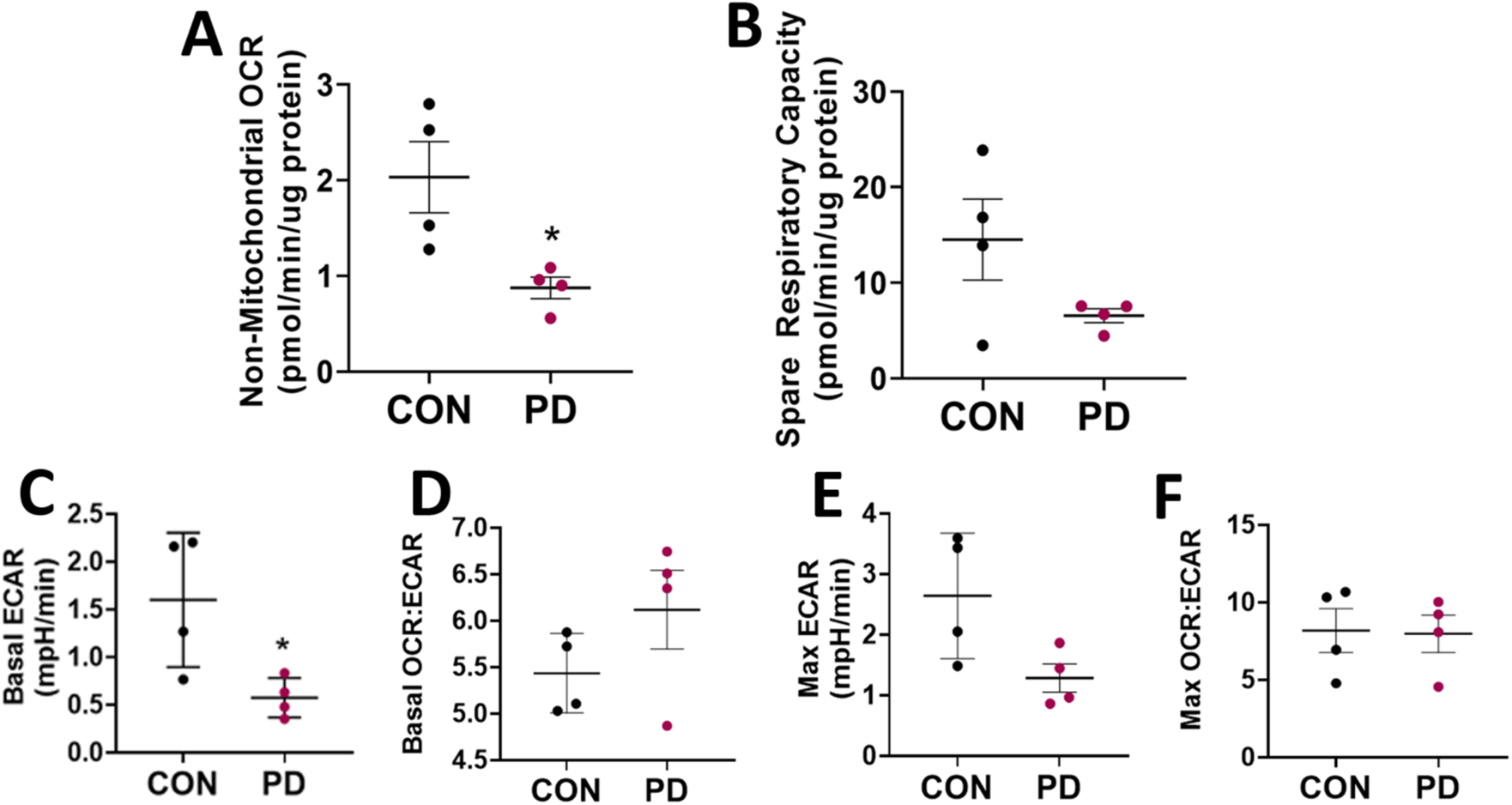
Seahorse Mito Stress analysis of iPSC-DAN. Seahorse mitostress test results relating to non-mitochondrial linked oxygen consumption and spare respiratory capacity from CON and PD cells is in (A, B). Basal ECAR and Maximal ECAR, and OCR/ECAR ratios are presented in (C-F). *p<0.05; Mean ± SEM, Unpaired t-tests, n=4 independent lines/group.

**Supplementary Figure 3:**
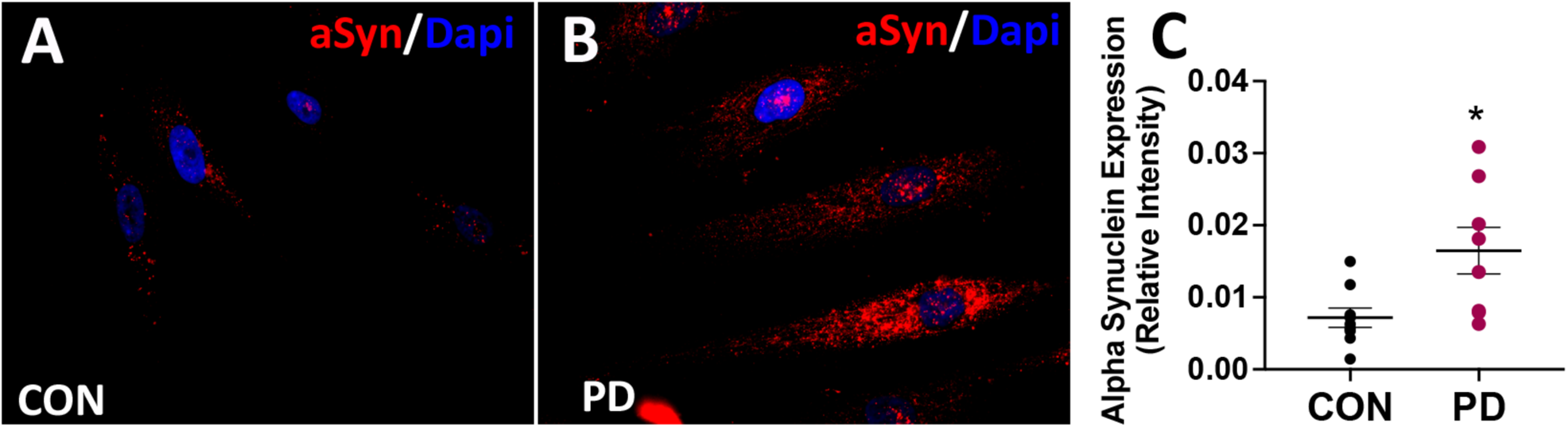
Comparative immunochemistry staining from control and PD fibroblasts is shown in (A, B). Results from quantification of the mean signal intensity is shown in (C). Scale bar - 50μM. *p<0.05; Mean ± SEM, Unpaired t-tests, n=8 independent lines/group.

**Supplementary Figure 4:**
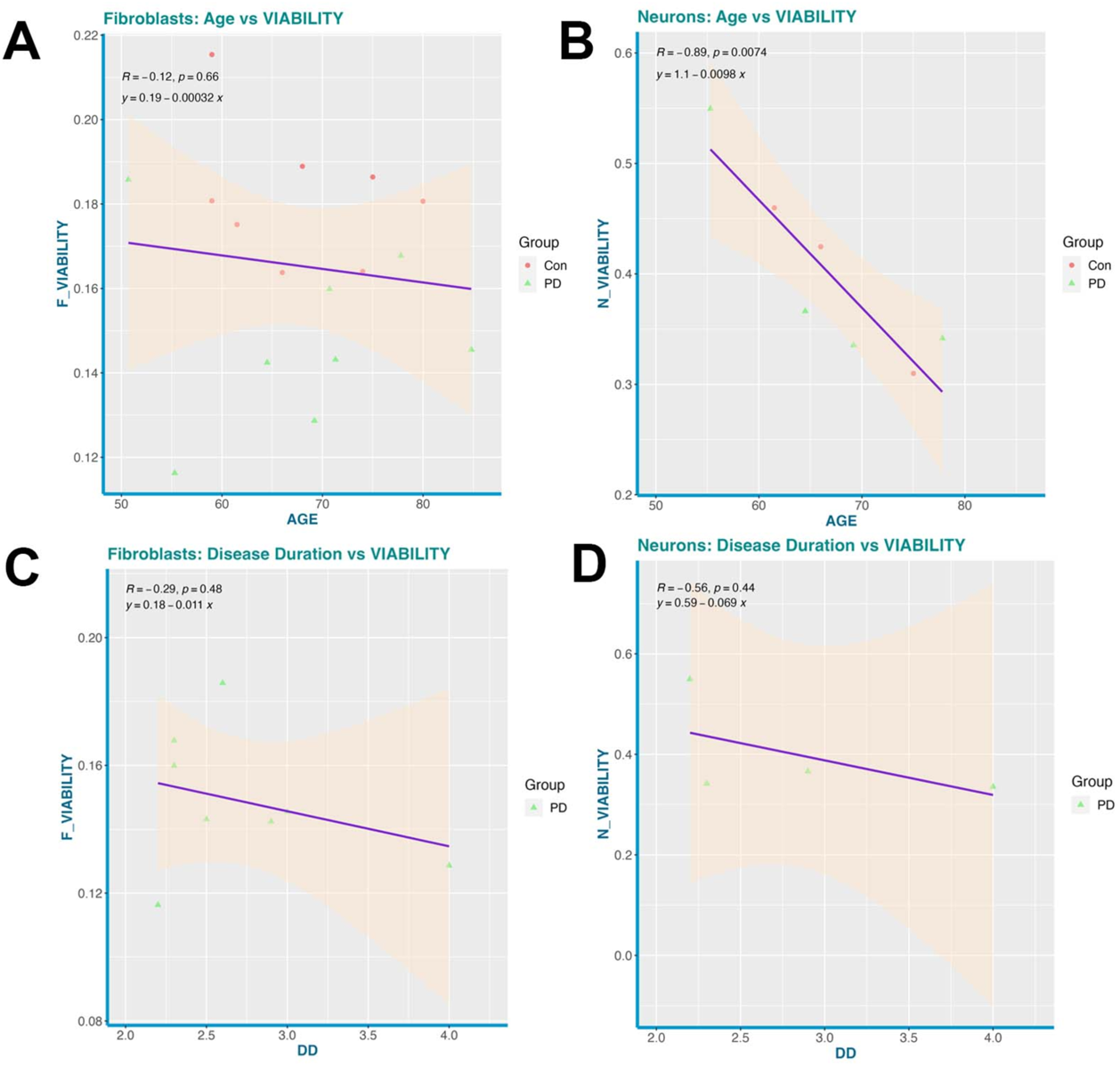
(A) and (B) show scatter plots capturing the linear relationship between age (age at sample collection) and viability (MTT assay) for neurons and fibroblasts, respectively. The linear relationship between disease duration (age at diagnosis to age at sample collection) and viability (MTT assay) is shown in (C) and (D).

## SUPPLEMENTARY METHODS

### iPSC culture and differentiation to midbrain DA neurons

Specifically, iPS cells were dissociated into single cells using StemPro Accutase (Gibco, ThermoFisher Scientific, Waltham MA) and plated at 1.8×10^5^–2.0×10^5^ cells per cm^2^ on Matrigel hESC-Qualified Matrix and allowed to expand until 75% confluent in mTeSR plus, typically overnight. Induction was then initiated by switching to KSR media: Knockout DMEM, 15% Knockout Serum Replacement, 2mM L-Glutamine, 1X Non-Essential Amino Acids, and 0.055mM β-mercaptoethanol (all from Invitrogen, Waltham MA), supplemented with LDN 193189 and SB 431542, which was designated as Day 0. In addition, on Days 1-2, SHH and Purmorphamine were added. CHIR was included in the supplemented KSR media starting on Days 3. On Days 5-10 a gradual shift in media was made and the neuralized cells were grown in varying concentrations of KSR media and N2 media [50% DMEM-F12, 50% Neurobasal, 2mM L-Glutamine, 1X N2 Supplement, 0.055mM β-mercaptoethanol, and 1X B-27 Supplement minus vitamin A (all from Invitrogen)]. Media concentrations were phased from 75%:25% (KSR:N2, Days 5-6), to 50:50 (Days 7-8), and to 25:75 (Days 9-10). In addition, SB 431542 was withdrawn on Day 5. As were SHH and Purmorphamine on Day 7. On day 11, media was changed to NB/B27 medium: 100% Neurobasal, 0.5mM L-Glutamine, 1X B-27 Supplement minus vitamin A and 50U/mL Penicillin-Streptomycin (all from Invitrogen). Days 11-12 NB/B27 media was supplemented with CHIR, BDNF, GDNF, TGF--b3, DAPT, AA, and cAMP in the concentrations mentioned above. From Day 13 onwards, CHIR was removed from the supplemented NB/B27 media and maturing DA neurons were fed every other day. Incremental increases in media volume were made as needed based on cell uptake.

On day 20, cells were pretreated for 30min with 10uM Y-27632 dihydrochloride (Selective ROCK Inhibitor: Tocris) and dissociated using TrypLE Express enzyme (Invitrogen). Cells were incubated at 37C in enzyme containing 10uM Y-27632 in 10minute increments, between which a 1mL serological pipet was used to gently triturate cultures into single cell suspensions. Line to line variability was noted in necessary enzyme exposure time and was closely monitored under a phase contrast microscope. Enzymatic activity was neutralized with the addition of 5 times the volume of Neurobasal media and centrifugation at 200 × G for 5minutes. Cells were replated under moderate density conditions (200×10^3^ cells per cm^2^) on dishes pre-coated with Poly-L-Ornithine/Fibronectin/Laminin [POFL: Poly-L-Ornithine (PO mw 30,000-70,000; 15μg/ml, Sigma-Aldrich), Fibronectin (4μg/ml, Corning), Laminin (6ug/mL, Sigma)]. Specifically, plates were coated with PO, made in Phosphate Buffered Saline without Calcium or Magnesium (PBS: Invitrogen), and incubated overnight at 37C. Plates were washed 3 times in PBS and coated with a combination of Fibronectin/Laminin, made in PBS and incubated overnight at 37C. POFL coated dishes were either stored at 4C or used immediately. Cells were cultured as described above until desired maturation stage for a given experiment, typically for 50-55 days.

## TABLE OF ANTIBODIES

| Antibody | Vendor | Catalog # | Host Species |
| --- | --- | --- | --- |
| anti-F-actin (Phalloidin) | ThermoFisher Scientific | A12379 | n/a |
| anti-ATPIF1 (5E2D7) | ThermoFisher Scientific | A21355 | Mouse monoclonal, IgG1 |
| anti-Tubulin Antibody, beta III isoform, C-terminus, clone TU-20 [Tuj1] | Millipore Sigma | MAB1637 | Mouse monoclonal, IgG1 |
| anti-Tyrosine Hydroxylase, clone LNC1 [TH] | Millipore Sigma | MAB318 | Mouse monoclonal, IgG1k |
| anti-FoxA2/HNF3β (D56D6) | Cell Signaling | 8186 | Rabbit monoclonal, IgG |
| anti-LMX1A | Millipore Sigma | ab10533 | Rabbit polyclonal, IgG |
| anti-Girk2 (Kir3.2) | Alomone | APC-006 | Rabbit polyclonal, IgG |
| anti-Map2 | ThermoFisher Scientific | PA110005 | Chicken polyclonal, IgY |
| anti-a-synuclein (Clone 42/a-Synuclein) | BD Biosciences | 610787 | Mouse monoclonal, IgG1 |
| anti-LC3 | Millipore Sigma | L7543 | Rabbit polyclonal |
| SQSTM1 monoclonal antibody (M01), clone 2C11 | Abnova | H00008878-M01 | Mouse monoclonal IgG2a kappa |

